# Comparative single-cell atlases reveal injury-driven tubal epithelial regeneration as a window for ovarian carcinoma initiation

**DOI:** 10.64898/2026.03.26.714557

**Authors:** Coulter Q. Ralston, Andrea Flesken-Nikitin, Dah-Jiun Fu, Christopher S. Ashe, Blaine A. Harlan, Md Mozammal Hossain, Derek K. Wang, Anna Yemelyanova, Elisa Schmoeckel, Andrew K. Godwin, Doris Mayr, Benjamin D. Cosgrove, Alexander Yu. Nikitin

## Abstract

High-grade serous carcinoma (HGSC), the most lethal form of ovarian cancer, preferentially originates in the tubal epithelium (TE) of the distal uterine tube (also known as Fallopian tube or oviduct). Mouse models are widely used to study how HGSC initiates in humans; however, the extent to which mouse and human uterine tubes are comparable remains unclear. Here, we conduct cross-species single-cell transcriptomic comparative analyses and organoid assay validations to reveal conserved differentiation trajectories from bipotent progenitors to secretory or ciliated cell fates. Regional analyses of both datasets reveal enriched injury repair features in the distal human TE, where mice lack such a trend. Experimentally inducing mechanical injury to the mouse TE yields significant expansion of pre-ciliated cells compared to uninjured counterparts. Furthermore, inactivation of *Trp53* and *Rb1,* whose pathways are commonly altered in HGSC, in regenerating pre-ciliated cells leads to rapid neoplastic transformation, implicating post-traumatic repair as a permissive window for malignant transformation. Together, our findings establish a comparative atlas of cell states between mice and humans, show that injury-associated regeneration may contribute to the known vulnerability of the fimbrial region, and raise potential concerns regarding procedures or conditions that mechanically perturb the tubal epithelium.

## Introduction

Ovarian cancer is the sixth leading cause of death for women in the United States ^1^. High-grade serous carcinoma (HGSC) is the most common and aggressive type of ovarian cancer, and most HGSC cases are detected in advanced stages where the 5-year survival rate is a bleak 32%; however, patients diagnosed with early stage HGSC improve their survival rate to 71% ^2,3^. To better treat patients with HGSC, it is critical to understand the susceptible cell states contributing to early oncogenesis.

Despite its namesake, HGSC most commonly originates in the distal tubal epithelium (TE) of the uterine tube, also known as the oviduct or Fallopian tube ^4^. A pivotal discovery for understanding early stages of HGSC came from the discovery of serous tubal intra-epithelial carcinomas (STIC) in prophylactic salpingo-oophorectomies from patients with *BRCA1* or *BRCA2* mutations ^5,6^. STIC lesions are most prevalent in the fimbriae specifically near junction areas at the mesothelial border in human uterine tubes ^6,7^. Although these precursors to HGSC provide evidence of early-stage disease characteristics such as *TP53* mutations, lesions present without symptoms and the origins of STIC development remain poorly understood ^8^.

In mice, *Pax8*+ cells were consistently found to self-renew and initiate HGSC upon inactivation of *Trp53* in tandem with other drivers ^9–12^. In fact, STICs are characterized as PAX8+ lesions in humans, thus alluding to a conserved cancer-prone cell state ^4^. However, *Pax8* is expressed in a broad spectrum of mouse tubal epithelial (TE) cell states, including stem cells, secretory cells, and pre-ciliated progenitor cells. In genetically modified mice, specific inactivation of driver genes in TE stem/progenitor cells marked by *Slc1a3* does not result in transformation ^12^. Instead, *Krt5*+ pre-ciliated cells are a rare cell state along the ciliogenic lineage that efficiently forms STIC lesions and HGSC upon inactivation of *Trp53* and *Rb1* ^12^.

Though the pre-ciliated cell state demands characterization in humans, previous single-cell RNA sequencing (scRNA-seq) studies fail to capture datasets expansive enough to combat patient batch variability and identify subtle epithelial cell states ^13–17^. As a result, rare cell state identification and definitive epithelial differentiation trajectories fail to reach a consensus. Additionally, it is imperative to understand how features of epithelial differentiation are conserved between mice and humans. This enables direct comparison between mouse models and humans for more accurate therapeutic discovery.

Regional factors that drive the fimbriae’s susceptibility to HGSC remain poorly defined. The classical hypothesis of ovarian carcinoma initiation (also known as incessant ovulation) is based on intensive postovulatory cell regeneration and proliferation of the ovarian surface epithelium (OSE), another putative tissue of HGSC origin ^18^. It has been suggested that similar regenerative mechanisms take place in the TE ^19,20^. However, no experimental evidence linking TE cell regeneration and carcinogenesis has been provided so far.

We assembled a comprehensive dataset integrating multiple publicly available datasets alongside our own to investigate rare cell states within the distal human TE. Using Self-Assembling Manifold mapping (SAMap ^21^), we generated a unified manifold to jointly analyze human and mouse TE cells. Through direct cell state comparison, we identified conserved epithelial lineage states between human and mouse data. We further examined regional variation within our human uterine tube dataset to identify features specific to distal TE. Collectively, our computational and experimental analyses implicate injury-induced regeneration as a key mechanism that may predispose the distal TE to neoplastic transformation.

## Results

### Atlas of cell types shared between the distal mouse and human uterine tube

Although the degree of coiling and relative size vary between species, the uterine tube in both humans and mice share paralleled regional distinctions ^12,22^. Ciliated cells are sparse within proximal regions, but prevalent in the distal regions of both species. Despite regionally distinct composition similarities, the uterine tubes of mice are surrounded by a protective bursa shielding the ovary from the peritoneal cavity, while humans rely on fimbriae to direct oocyte capture while brushing along the ovary open to the peritoneal cavity (Figure 1a-b). To compare the cellular features of mouse and human uterine tubes, we analyzed single-cell RNA-sequencing (scRNA-seq) samples from multiple sources. The samples from mice were all estrous adults between 8-12 weeks with the samples split into proximal and distal regions (Supplementary Figure 1a). The human samples were pooled among four sources to account for a wide range of regional distinctions and ages (Supplementary Figure 1b). To consistently organize these datasets, we processed the raw sequencing data from each source, followed standard preprocessing by removing low quality cells, and used harmony for batch correction (Supplementary Figures 1c-e and 2).

**Figure 1.**
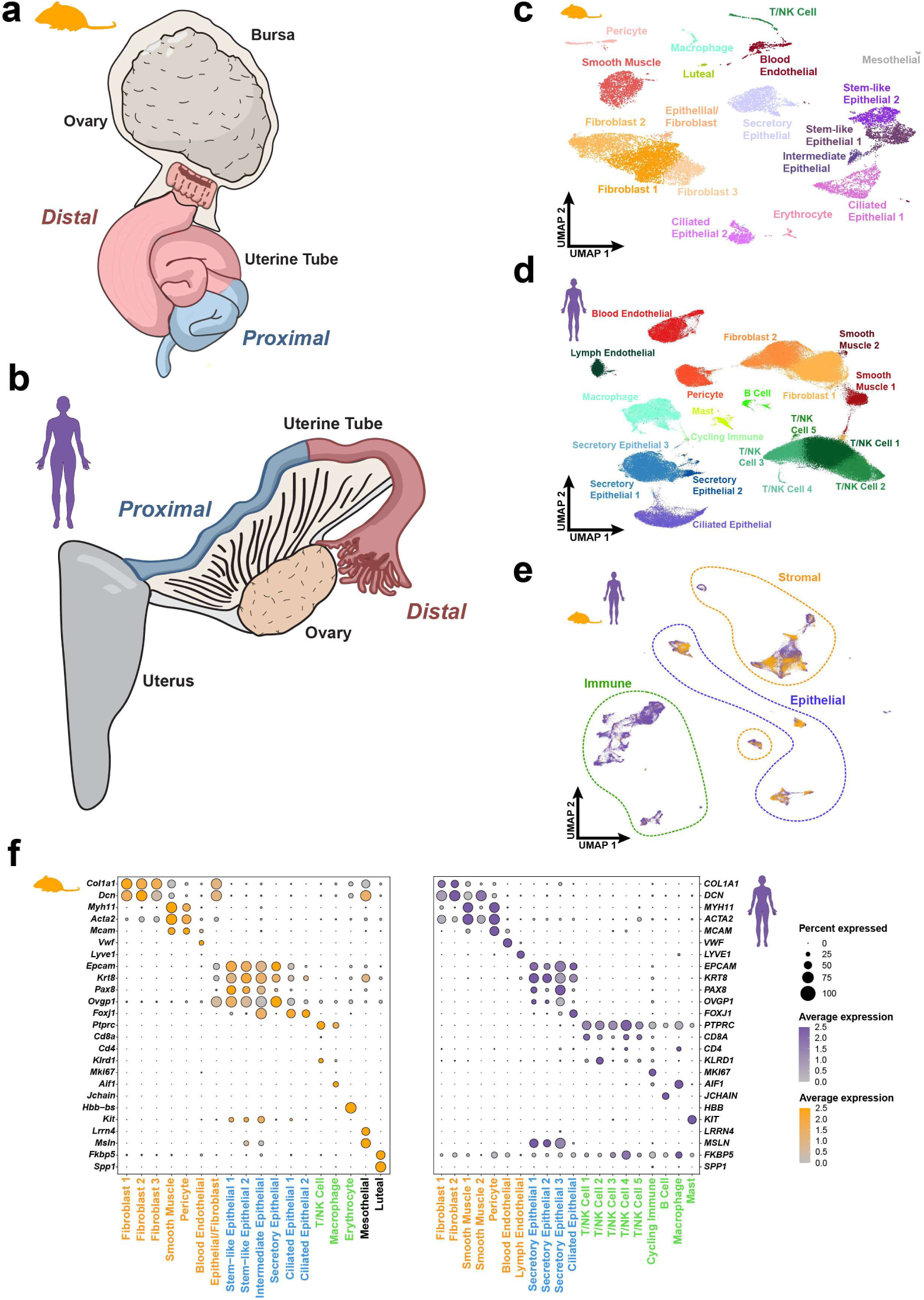
Cell types are conserved between mice and humans in the distal uterine tube. (**a-b**) Diagram of uterine tubes found in the mouse (oviduct, a) and human (Fallopian tube, b). The proximal region and distal region are generally conserved with greater ciliated cell abundance in the distal regions. (**c-d**) UMAP visualization of the cell types identified in a pool of 16,583 distal mouse uterine tube cells (c) and 226,536 human uterine tube cells (d). (**e**) SAMap representation of combined mouse and human uterine tube single-cell transcriptomes. (**f**) Dot plot representation of conserved cell type gene homology shared between cell types in humans and mice.

**Figure 2.**
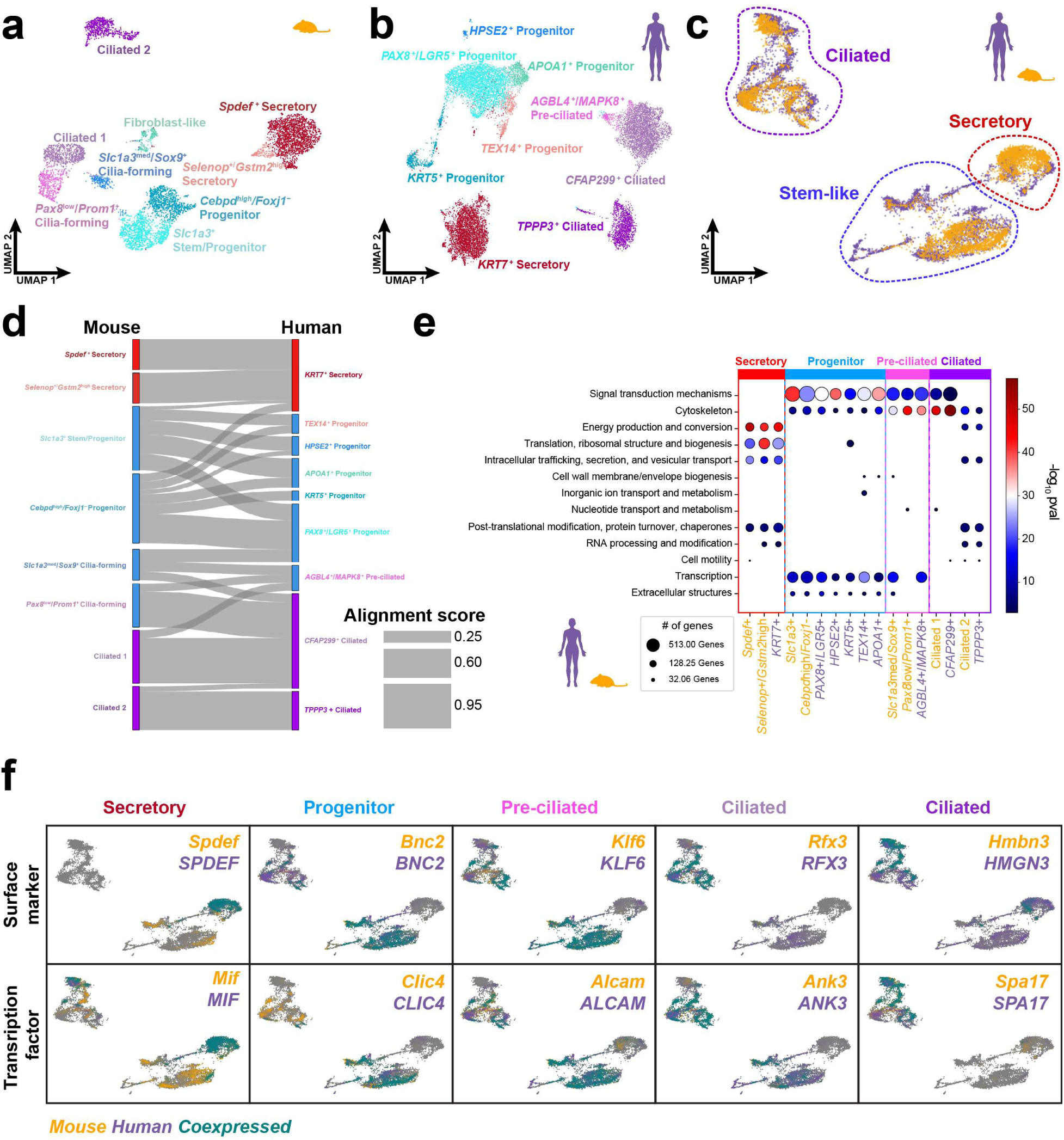
Epithelial cell states are conserved in mice and humans. (**a-b**) UMAP visualizations of the distal mouse epithelial cells (a) and the human fimbria epithelial cells (b). (**c**) Combined human and mouse distal tubal epithelial object with conserved major epithelial cell types labeled. (d) Cell states conservation based on mapped cell state clusters in humans and mice visualized in a Sankey plot. (**e**) Functional annotations of eukaryotic orthologous groups across epithelial cell states in both species. (**f**) Feature plots of homologous cell surface marker and transcription factor RNA transcripts identified in mouse and human epithelial cell states.

In representative samples from 31 adult mice and 23 human patients, we identified 16,583 and 226,536 high quality cells from distal uterine tubes respectively, which then were categorized into 18 clusters representing cell types in mice and 20 clusters in humans (Figure 1c-d). Although the human samples all had different collection and preservation methods, major cell types were well distributed in all samples encouraging further unbiased exploration of the dataset (Supplementary Figure 3). To map the extent to which the identified cell types from mice and humans are conserved, we employed SAMap to identify homologous cell types and shared gene expression patterns across species ^21^. Epithelial, stromal, and immune cell populations exhibited conserved representation within the distal uterine tube across both (Figure 1e). However, there are notable differences including increased immune cell abundance, lymphatic endothelial cells, and the presence of mast cells in human datasets but not in mice (Figure 1f).

### Distal epithelial cell state conservation between mice and humans

While revealing overall cell type contributions are important, the epithelial cell compartment is especially crucial for elucidating its role in the origin of HGSC. We applied the same approach as stated prior to apply SAMap on epithelial subsets. We subset 6,273 distal epithelial cells from mice, which were categorized into 8 cell states after removing the fibroblast-like cluster due to doublet concerns (Figure 2a). In humans, 62,703 distal epithelial cells were subset and separated into 9 cell states (Figure 2b). Epithelial cell states in human and mouse subsets were characterized by their most highly differentially expressed feature(s). Human distal epithelial progenitor populations were the most diverse based on their variety of unique features. However, both distal epithelial subsets shared similar features of stem/progenitor, secretory, and ciliated cell states. After applying SAMap to the epithelial subsets, ciliated, secretory, and stem-like bins were identified within the mapped manifold (Figure 2c). Ciliated cells and secretory cells showed the largest alignment scores, while stem/progenitor clusters were difficult to align (Figure 2d).

A better understanding of cell state conservation relied on functional annotations of eukaryotic orthologous groups (KOG ^23^). KOG analysis revealed a shared emphasis of signal transduction and cytoskeleton features on stem/progenitor and pre-ciliated cell states of both species, while energy production and conversion were upregulated in differentiated ciliated and secretory cells (Figure 2e). Furthermore, we identified conserved markers with experimental relevance, such as homologous cell surface markers and transcription factors for each conserved cell state (Figure 2f). Among those, progenitors shared overall expression of markers (*BNC2* and *CLIC4*) regardless of species or subcluster-specific features.

While conservation of homologous cell states was essential to translating mouse and human genes, we sought out to establish a reference that also includes species-specific expression of features. We first identified the most distinguishing markers for each epithelial cells state in humans and mice. Then, we calculated correlation scores between defining cell state markers and their respective orthologs across species. From there, we identified the most conserved, human-specific, and mouse-specific markers for each cell state (Figure 3). The distal epithelial conservation reference reflects the Sankey plot (Figure 2d) with abundant conserved features in ciliated and secretory cells, while progenitor populations had more specific markers. Together, the conservation reference for epithelial cell states can serve as a basis for further exploration between species.

**Figure 3.**
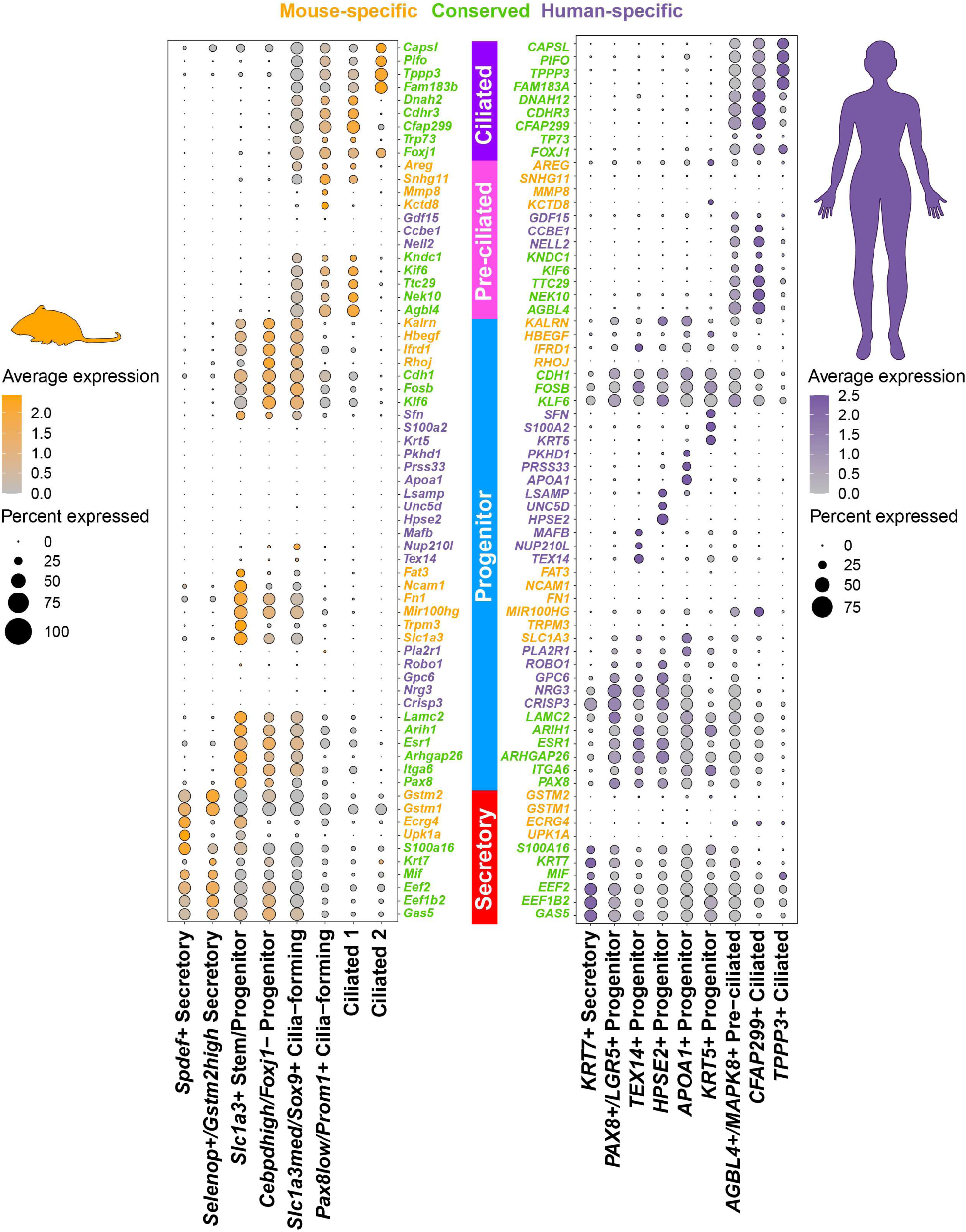
Distal epithelial conservation reference for features of distal TE cell states. Conserved (green), mouse-specific (yellow), and human-specific (purple) feature characterizing the distal epithelial cells in mouse and human uterine tubes.

### Establishing distal epithelial lineage dynamics shared between mice and humans

Mapping mouse to human uterine tube sequencing data enabled the characterization of human epithelial cell state lineages with supported mouse lineage tracing data. The epithelial lineage hierarchy of uterine tubes has been well defined in mice ^12^. However, limited resolution across individual datasets has led to conflicting reports regarding epithelial lineage trajectories in human uterine tubes. To assess lineage conservation between species, we utilized Monocle3 pseudotime trajectory analysis applied over PHATE embeddings ^24–27^ to visualize epithelial differentiation trajectories in mouse and human datasets (Figure 4a-f). Using established markers of mouse epithelial differentiation, we leveraged homologous gene pairs identified by SAMap to construct a pseudotime-binned trajectory analysis, thereby enabling direct comparison of conserved features of epithelial differentiation across species (Figure 4g). Comparatively, analysis revealed that the human trajectory exhibited less robust differentiation along the secretory lineage, as evidenced by fewer pseudotime bins with high *SPDEF* expression relative to *Spdef* in the mouse trajectory. In humans, *KRT7* marked the secretory lineage, with increased average expression in left-most pseudotime bins; however, its expression in mice extended across both epithelial lineages. Progenitor-associated markers (*SLC1A3*, *ITGA6*, *PAX8*) were detected in human progenitor bins, though with reduced specificity compared to mouse trajectories. While the pre-ciliated cell marker *Krt5* was not conserved in human trajectories, other markers of ciliogenesis demonstrated stronger conservation and specificity. Notably, both species exhibited a transition from *TP53* to *TP73* expression during early stages of ciliogenesis, followed by enrichment of *TPPP3* and *FAM183B* in later stages, consistent with conserved programs of ciliated cell differentiation.

**Figure 4.**
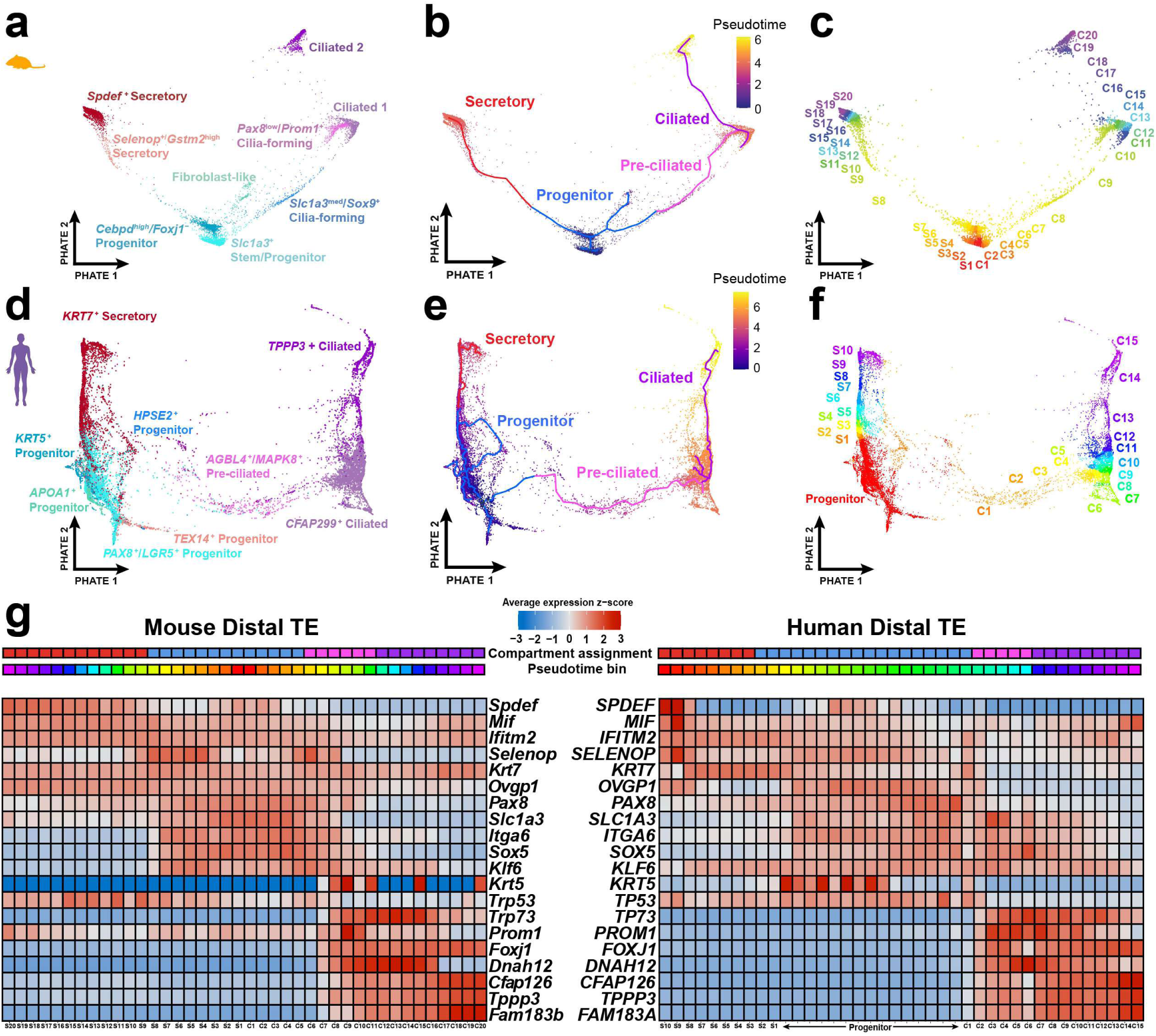
Epithelial pseudotime trajectories project shared mechanisms regulating homeostatic tissue regeneration in mice and humans. (**a-c**) Embedding of distal mouse tubal epithelial single-cell transcriptomes using PHATE (a). Where monocle3 pseudotime trajectory inference overlays a path for progenitor, secretory, pre-ciliated, and ciliated cell compartment assignments (b). This embedding was then split into 40 equally sized bins across the pseudotime trajectory (c). (**d-f**) Embeddings of human tubal epithelial single-cell transcriptomes were analyzed the same as the mouse for the PHATE visualization (d), monolce3 trajectory inference (e), and bin assignment (f). (**g**) Shared expression trends across pseudotime bins compared across species. The average expression of features calculated across equally spaced bins with supporting data from cell state compartment assignment.

### SAMap predicts analogous gene pairs for epithelial stem/progenitor cell conservation in humans and mice

SAMap supports the identification of correlated genes expressed across mapped cell types. While differentiated epithelial cell states were highly conserved, stem/progenitor cell state mappings and specific markers were ambiguous within our pseudotime trajectory analysis. To investigate whether we can employ SAMap to identify previously undescribed genes without the bias of gene homology, we used correlation scores and blast scores of gene pairs to identify analogous cell surface marker genes from mouse *Slc1a3*+ stem/progenitor cells (Figure 5a). To further refine the list of genes, we filtered for the highest specificity according to pseudotime lineage trajectories so that progenitor state diversity in humans was represented (Figure 5b). Among the top 10 candidate genes were *ROBO1* and *PLA2R1*, which had not been previously described as a stem-like marker in the human TE. Both *ROBO1* and *PLA2R1* were specifically expressed in progenitor pseudotime bins, while secretory marker *KRT7* and ciliated marker *FOXJ1* represented their respective trajectories (Figure 5c).

**Figure 5.**
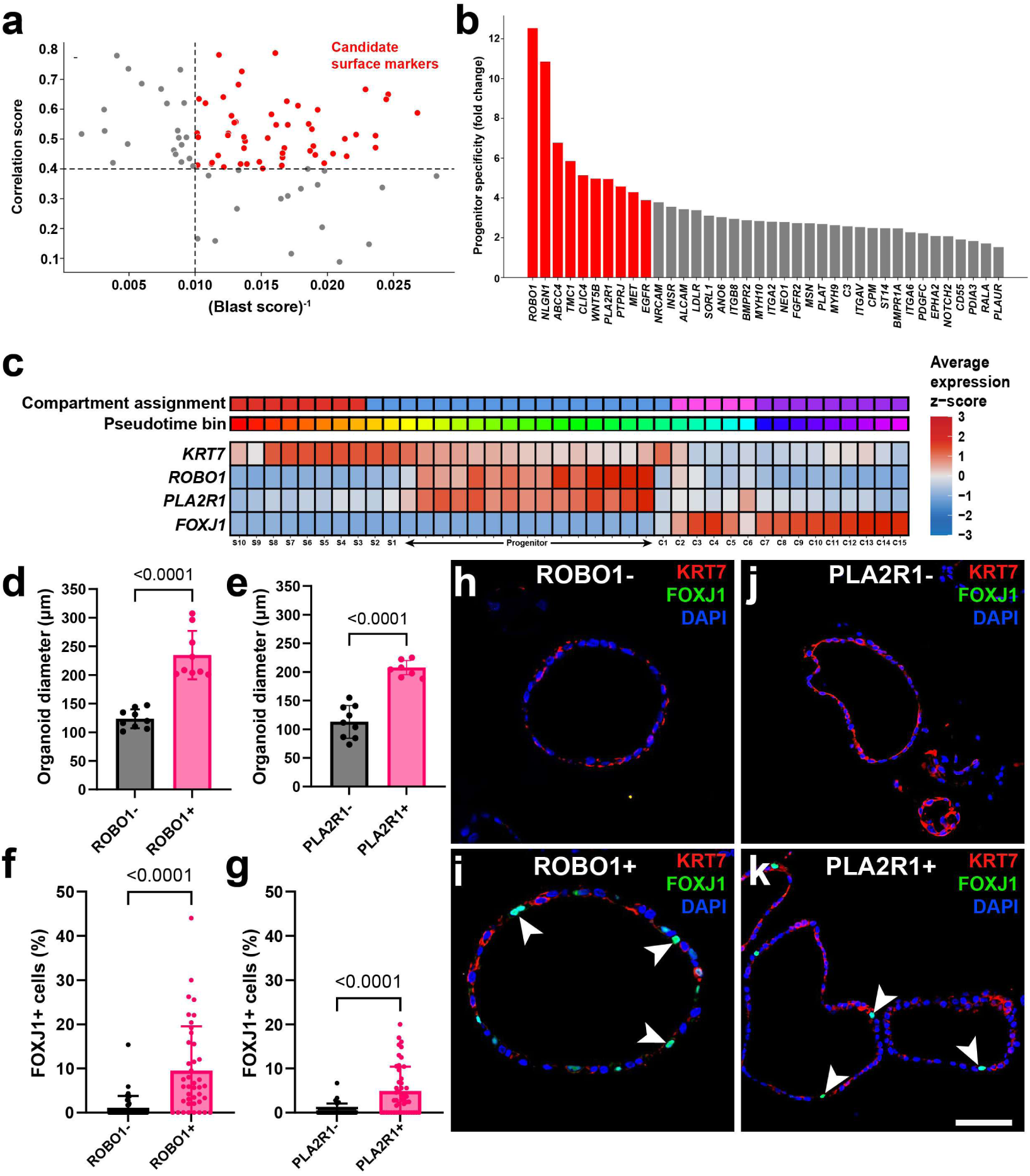
Analogous stem/progenitor cell markers predicted by SAMap select for organoid-forming cells capable of differentiation. (**a-b**) Selection criteria for stem/progenitor cell gene pairs identified by SAMap. The top gene pairs encoding for surface markers between stem/progenitor populations in both species were filtered for the highest correlation scores and the lowest blast scores (a). The filtered candidate surface marker list ranked in the order of pseudotime lineage specificity (b). (**c**) Average z-score expression trends across pseudotime bins for features characterizing the distal human TE lineage hierarchy. (**d-e**) Organoid diameter measurements between positive bound and unbound fractions from ROBO1+ (d) and PLA2R1+ (**e**) cells isolated from human uterine epithelial cells by MACS. (**f-g**) Quantification of FOXJ1 positivity reflecting enrichment of ciliated cells within ROBO1+ (f) and PLA2R1+ (g) organoids. (**h-k**) Representative images for organoids formed from ROBO1- (h), ROBO1+ (i) PLA2R1- (j), PLA2R1+ (k) epithelial cells stained for secretory marker KRT7 (red), ciliated marker FOXJ1 (green), and DAPI (blue). Scale bar 50 μm (h-i) and 70 μm (j-k). Statistics by Welch’s t-tests. Bars denote SD.

To validate SAMap’s ability to predict analogous gene pairs, we performed magnetic-activated cell sorting (MACS) to split human uterine tube epithelial cell pools into marker-bound and unbound groups. Organoids derived from both ROBO1+ and PLA2R1+ cells formed significantly larger organoids than the unbound epithelial cells (Figure 5d-e). All organoids in selected or unbound conditions expressed secretory marker KRT7 by immunofluorescent staining (Figure 5h-k). However, the organoids from ROBO1+ and PLA2R1+ cells expressed significantly greater expression of ciliated marker FOXJ1 (Figure 5f-g). These experiments suggested that both ROBO1 and PLA2R1 are markers of stem/progenitor epithelial cells in humans with the capacity to differentiate into ciliated progeny, and SAMap can identify analogous gene pairs in cases were mapped homologous genes are sparse.

### Upregulated features of the human fimbriae suggest a link between mechanical damage and cancer susceptibility

After establishing a reference for conservation between human and mouse TE lineage hierarchies, we explored the human dataset to identify regional differences that may be associated with HGSC pathogenesis. Regional analyses uncovered fimbriae-specific features to interrogate why this region preferentially forms STIC lesions. In the human dataset, the relative abundance of immune cell populations increased distally, while the mouse dataset showed opposite results with the proximal region being the most abundant in immune populations (Supplementary Figure 4a-b). Enriched markers for fimbriae fibroblast and macrophage populations revealed injury-associated genes, such as the case of *CCL20* in macrophages; nevertheless, this regionally specific marker was not translatable to mice (Supplementary Figure 4c-f).

Notably, fibroblasts from the human fimbriae exhibited increased *POSTN* expression, a marker associated with chronic wound repair (Figure 6a-b). In contrast, mouse fibroblast datasets did not demonstrate regionally-specific *Postn* expression, suggesting comparatively reduced activation of wound repair pathways (Figure 6c). These findings may reflect anatomical differences between species. In mice, the ovarian bursa provides relative rigidity to the distal uterine tube and likely protects it from mechanical injury. In humans, by contrast, the distal uterine tube opens into the peritoneal cavity, and the fimbriae sweep and brush over the ovary. This persistent epithelial perturbation may drive ongoing tissue damage and compensatory regeneration responses in the human distal uterine tube.

**Figure 6.**
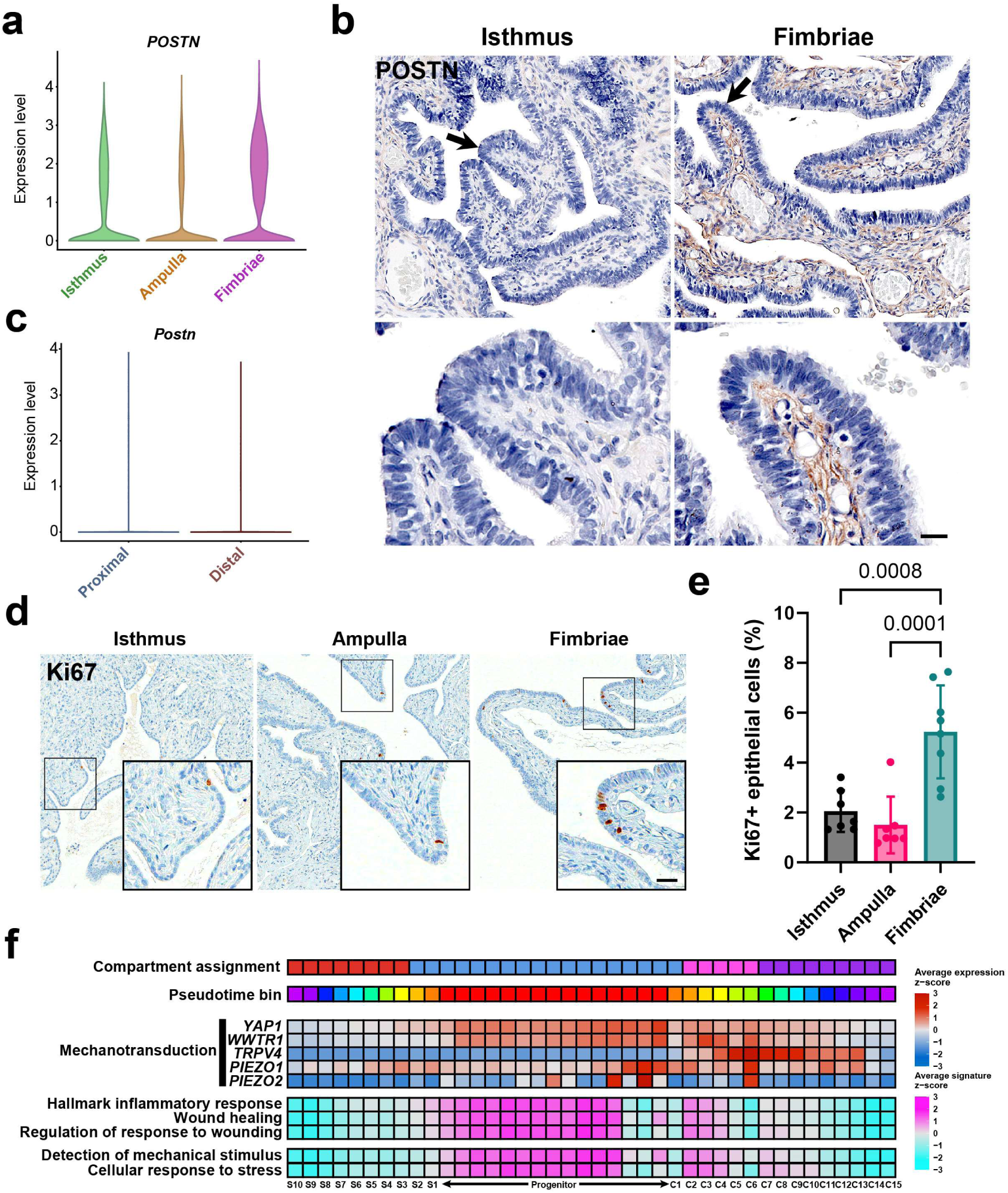
Features of the human fimbriae suggest mechanical damage as a risk factor for HGSC initiation. (**a**) Violin plot of regional expression of *POSTN* in the human uterine tube. (**b**) Immunostaining for periostin (brown) in human isthmus and fimbriae, Elite ABC method, hematoxylin counterstaining (right column). Scale bar, 50 μm top row, 17 μm bottom row. (**c**) Violin plot of regional expression of *Postn* in the mouse uterine tube. (**d**) Immunostaining for Ki67 (brown) in epithelial cells from isthmus, ampulla, and fimbriae regions of the human uterine tube. Scale bar, 60 μm whole images, 24 μm insets. (**e**) Quantification of Ki67+ epithelial by region. One way Anova, Tukey multiple comparisons post hoc tests. Bars denote SD. (**f**) Human pseudotime binned feature trends for mechanotransduction-associated genes and gene signatures associated with stress and healing. Expression of individual genes and signatures of gene sets were calculated for their normalized z-score expression across pseudotime bins.

Consistent with increased demand for regeneration, frequency of transitional pre-ciliated cells was higher in the human fimbriae compared to the ampulla (Supplementary Figure 4g). The fimbriae also demonstrated an increase in epithelial turnover, as marked by more than two-fold greater abundance of Ki67+ cells in the fimbriae compared to the ampulla and isthmus in human samples (Figure 6d-e). Moreover, fimbriae epithelial lineage trajectories revealed an enrichment for genes involved with mechanotransduction specific to the pre-ciliated populations (Figure 6f). The pre-ciliated and progenitor populations also expressed upregulated feature signatures relating to wound healing and detection of mechanical stress. Together, these findings implicate pre-ciliated cell population as a key mediator of post-traumatic regeneration, a process that appears to be selectively active in the exposed, fimbriated regions of the human uterine tube.

### Inducing mechanical injury in mice stimulates pre-ciliated cell expansion and exacerbates HGSC initiation

Previously we reported that *Krt5*+ pre-ciliated cells were previously found to initiate HGSC after inactivation of *Trp53* and *Rb1* in mice ^12^. To investigate how wounding affects pre-ciliated cell populations, we developed a microsurgical technique to damage the distal mouse uterine tube. Injuries to K5-CreERT2 Ai9 mice at the same time as tamoxifen treatment resulted in significant expansion of *Krt5*+ progeny compared to uninjured contralateral uterine tubes (Figure 7a-d). Injuring TE K5-CreERT2 *Trp53^loxP/loxP^ Rb1^loxP/loxP^* Ai9 mice resulted in formation of dysplastic lesions formed significantly earlier in traumatized TE as compared to intact TE of the same mouse (26 days vs. 120 days after induction, with vs. without injury (Fischer’s exact P=0.0003; Figure 7e-n and Supplementary Table 1). Such lesions consisted of cells with pronounced atypia, loss of cell polarity, and high proliferative index typical for STICs (Figure. 7c). They also presented with expression of PAX8, Wilms tumor 1 (WT1) and P16. Trauma induction also significantly increased the frequency of carcinomas similar to human HGSC (100% vs 44%, with vs. without injury, n=18, Fischer’s exact P=0. 0116). Such carcinomas showed slit-like and solid patterns of growth, expression of PAX8, WT1, P16, high levels of proliferation and stromal invasion (Supplementary Figure 5 and Supplementary Table 1). Together, mechanical damage of the TE has led to the expansion of pre-ciliated cells and HGSC formation, thereby directly linking TE regeneration and carcinogenesis.

**Figure 7.**
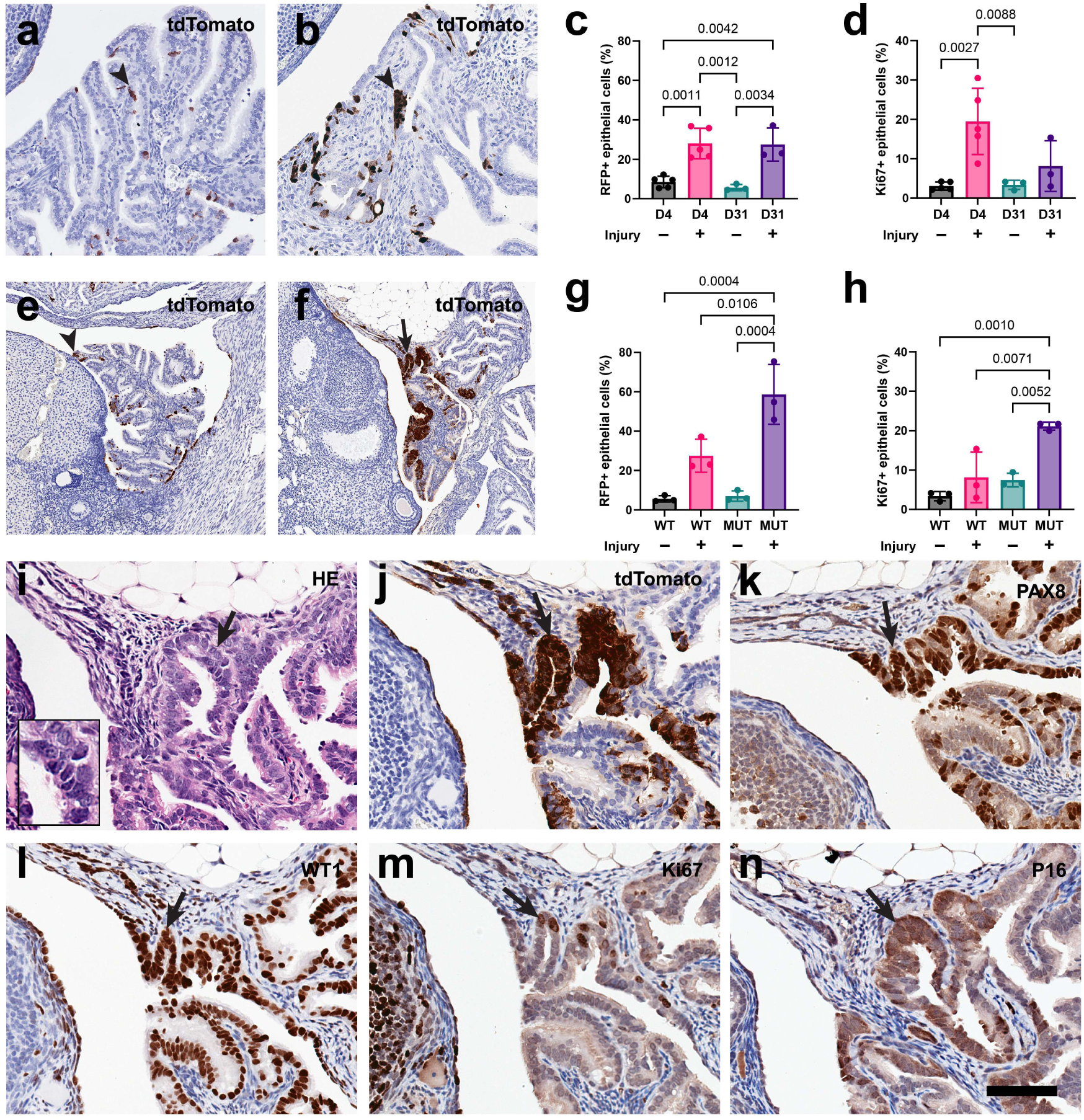
Mechanical injury in mice leads to expansion of pre-ciliated cells and acceleration of HGSC formation. (**a-b**) tdTomato expression (brown) in contralateral uninjured (a) and injured (b) uterine tubes of K5-CreERT2 Ai9 mice 4 days after tamoxifen administration. (**c-d**) Quantifications of epithelial cells expressing RFP (c) and Ki67 (d) from uninjured and injured uterine tubes between day 4 and 31 in K5-CreERT2 Ai9 mice (WT). (**e-f**) tdTomato expression (brown) in contralateral uninjured (e) and injured (f) uterine tubes of K5-CreERT2 *Trp53^loxP/loxP^ Rb1^loxP/loxP^* Ai9 mice 35 days after tamoxifen administration. (**g-h**) Quantifications of epithelial cells expressing RFP (c) and Ki67 (d) from uninjured and injured uterine tubes from WT and K5-CreERT2 *Trp53^loxP/loxP^ Rb1^loxP/loxP^* Ai9 mice (MUT) 31 days after tamoxifen treatment. (**i**-**n**) Larger magnification of dysplastic lesion shown in i (arrow) and characterized by cellular atypia, loss of cell polarity, and high proliferative index typical for STICs (inset). Staining with hematoxylin and eosin (HE, i), tdTomato (j), PAX8 (k), WT1 (l), Ki67 (m), and p16 (n). (a-b, ef, i-n) Elite ABC method, hematoxylin counterstaining. Regular (arrowhead) and atypical (arrow) TE cells. Scale bar, 120 µm (a, b), 240 µm (c, d), 60 µm (e-j) and 30 µm (e, inset). Statistics by one way Anova, Tukey multiple comparisons post hoc tests. Bars denote SD.

## Discussion

Our single-cell transcriptomic reference connecting mouse and human uterine tube scRNA-seq datasets elucidated similarities and differences among all cell types. Between humans and mice, we found conservation of general fibroblast, macrophage, and epithelial cell type markers. However, mice lacked a notable abundance of immune populations, whereas human uterine tubes included T/NK cells, mast cells, and B cells. Further analysis of specific features of human datasets revealed divergent features of shared cell types such as *CCL20* expression in macrophage and *POSTN* in fibroblasts. Understanding differences between mouse and human cell types can better inform mouse model experiments to better recapitulate human physiology.

Specific investigation into epithelial cell trajectory conservation revealed which features of human and mouse epithelia aligned. Imprinting conserved genes investigated in mice enabled the characterization of human epithelial cells which previously lacked sufficient validation. Within mouse and human datasets, we established a reference of shared and exclusive genes in each epithelial cell state. Using computational lineage trajectory inference, conserved features in both humans and mice suggested a stem/progenitor cell state capable of secretory and ciliated cell differentiation trajectories. These data are supported by previous lineage tracing studies that identify *ITGA6* (CD49f), *Pax8*, and *Slc1a3* as progenitor markers for the fallopian tube, albeit not as specific in human trajectories ^9,12,19^.

To improve consistency in the prediction of specific cell states, we leveraged SAMap to identify analogous genes that are functionally similar but evolutionary distinct genes across in species. Focusing on stem/progenitor populations, we prioritized candidate markers based on cross-species correlation, cell type, specificity, and lack of sequence homology. Among the top candidates identified in humans *ROBO1* and *PLA2R1* emerged as robust markers of stem/progenitor TE cells. These populations demonstrated enhanced functional capacity, forming larger organoids and giving rise to both KRT7+ secretory and FOXJ1+ ciliated cell lineages. Collectively, these findings broad the repertoire of markers for identifying and characterizing uterine tube epithelial cell states and provided a framework for improved cross-species annotation of stem/progenitor populations.

One notable advantage over previous studies is the resolution in which we characterize the human uterine tube. Previous studies were based on a limited number of diverse patient samples in individual datasets. As a result, studies advocated for contradictory views of the human TE lineage hierarchy ^13–16^. We combined previously published datasets and included our own to gain a better picture of uterine tube dynamics over regional distinctions. This facilitated more controlled comparative analyses across species.

Regional assessment uncovered features of mechanical injury enriched in the human fimbriae. Others provide additional evidence of this with increased apoptotic features being expressed in the fimbriae ^17^. Many studies have hypothesized that DNA damage caused by follicular fluid may impact HGSC risk by increasing inflammation ^20,28–31^. However, these reports alone have yet to definitively identify follicular fluid as a true driver of HGSC progression. Specifically, ovulation did not result in proliferation of TE cells in mouse models, nor was this the case in organotypic and cell culture from baboon fimbriae ^28^. Although proliferation was not observed, increased DNA damage to the TE and macrophage infiltration occurred in mice with increased ovulation ^28^. This observation suggests that other local stimuli exclusive to the fimbriae may be responsible for the regional bias of STIC and HGSC progression.

We find that *POSTN* gene expression is significantly upregulated in the fimbriae fibroblasts of humans but not in the distal region of mice. Periostin is a known marker of wound repair, where its role in healing is conserved across multiple tissues^32^.

Furthermore, *POSTN* expression was reported to be upregulated in lung chronic injury models compared to uninjured control groups^33^. These findings support the hypothesis that the protective bursa encapsulating the uterine tube in mice shields it from continuous mechanical damage, while the freely moving fimbriae in humans are exposed to constant mechanical trauma, thus resulting in periostin upregulation.

To evaluate whether mechanical stresses could induce TE proliferation, we first investigated which cell states were associated with mechanotransduction and wound repair. Among the upregulated cell states were stem/progenitors and pre-ciliated cells. Pre-ciliated cells exclusively expressed *TRPV4*, a mechanosensor responsive to osmotic and mechanical stimuli. TRPV4 has previously been identified for its upregulation during viscous loading of fallopian tube epithelial primary cell cultures ^34^. In addition to *TRPV4*, other mechanotransduction effectors such as piezo channels, YAP, and TAZ were all upregulated in human pre-ciliated cells ^35^. The shared expression of these features suggests that pre-ciliated cells are capable of mechanotransduction in response to mechanical stimuli. Additionally, these mechanically-dependent features can lead to increased cancer cell survival and migration, thus implicating pre-ciliated cells as a conserved cancer-prone cell state ^36^.

Upon mechanical injury to mouse uterine tubes, pre-ciliated cells were confirmed to expand rapidly compared to uninjured controls. This implies that pre-ciliated cells are implicated in post-traumatic regeneration and further suggests that mechanical damage could serve as a risk factor for HGSC. In fact, mechanical injury to mouse uterine tubes resulted in rapidly forming lesions that appear as soon as 26 days after treatment. The increased formation rate of HGSC in mechanically damaged mouse uterine tubes informs surgical techniques where only partial removal of the uterine tube is completed. Recent characterization of the fimbria ovarica, a microanatomical structure connecting the human fimbria to the ovary, could be implicated as it may be retained after prophylactic salpingectomy procedures ^37^. Altogether, we propose that pre-ciliated cells are a therapeutic target for early transformation events and mechanical damage during chronic damage and prophylactic removal procedures intensifies HGSC progression.

Although this study focused on epithelial cell lineages and their contribution to HGSC initiation, the dataset provides a broader resource for conserved cell types with applications in disease modeling, therapeutic target identification, and the study of shared developmental programs. Beyond the uterine tube, our findings highlight the utility of SAMap-based identification of analogous gene pairs as a generalizable framework for cross species tissue characterization. Applications of this approach across additional tissues and model systems may enable more precise annotation of cell states and accelerate the discovery of conserved, functionally relevant therapeutic targets.

## Materials and Methods

### Human patient and tissue acquisition

De-identified clinical samples used for single-cell RNA-sequencing (scRNAseq) and TE organoid preparation were obtained from the Biospecimen Repository Core Facility (BRCF) at the University of Kansas Medical Center (KUMC) together with relevant clinical information. The use of human samples was approved by the Institutional Review Board under the HSC #5929 at KUMC. Human fallopian tube tissue samples obtained from women who provided informed consent were de-identified using OpenSpecimen prior to distribution and analysis. Unless otherwise indicated, all sample handling and experimental procedures were conducted in a biosafety level 2 (BSL-2) cabinet. Distal (fimbriae and ampulla) and proximal (isthmus) regions of the uterine tube were collected on ice from standard surgical procedures for benign gynecological disease. The surgical tissue samples were examined by a surgical pathologist (maintaining the orientation and laterality of the sample) and only anatomically normal uterine tubes were used. Samples were frozen in Cryopreservation Medium M-TE-N1 ^38^, shipped on dry ice, and kept for long term storage in liquid nitrogen.

De-identified sectioned slides of uterine tubes collected from patients with and without HGSC were prepared at the Institute of Pathology, Ludwig Maximilians University, Munich, Germany and Weill Cornell Medicine (Cornell IRB 0005546 and Weill Cornell Medicine IRB protocol 23-07026303).

### Single-cell RNA-sequencing library preparation

For the mouse TE (mTE) single-cell RNA-sequencing (scRNAseq) library data sets we utilized our previously reported transcriptomes ^12^.

For human TE scRNAseq analysis we isolated primary cells from cryopreserved patient tissue samples. Three primary cell isolation experiments D33, D34, and D35 were performed with a collagenase type I digestion buffer ^39^. Briefly, one vial of cryopreserved tissue was thawed for 2-3 minutes in a 37°C water-bath. Next the tissue was washed three times in 15 ml Wash Buffer (PBS, 1x Phosphate buffered saline pH 7.4/Penicillin, Streptomycin 100 IU ml^-1^, 100 µg ml^-1^; Corning 21-031-CV; PS, Corning 30-002-Cl). Leaving the tissue in the third Wash Buffer, using scalpel blades, tubes were opened longitudinally to expose the mucosa folds, and connective vascular tissue was removed. Afterwards, the tissue was transferred into 10 ml Digestion Buffer (0.5 mg ml^-1^ collagenase type I (Thermo Fisher 17100-017) in Advanced DMEM/F12 (ADF, Thermo Fisher 12634-028), containing 12 mM HEPES (HEPES, 1M, Thermo Fisher 15630-080 ) and 3 mM calcium chloride (CaCl_2_, 1 M, Bioworld 40320005-1) for 45 minutes in a 37°C water-bath with gentle shaking every 10 minutes. After the incubation the tissue was place with 2 ml Digestion Buffer on a 10 cm dish lid and minced in 2 – 4 mm pieces, combined with the remaining 8 ml, and mixed 20 times. The tissue cell pellet was collected by centrifugation, 300xg for 3.5 minutes at 25°C, and suspended with 30 ml A-TE-2D medium (5% fetal bovine serum (FBS, Sigma Aldrich 12306C) in ADF supplemented with 12 mM HEPES, 1% L-GlutaMaX-I (Thermo Fisher 35050-079), PS same concentration as Wash Buffer, 10 µM Rho kinase inhibitor (Y-27632, ROCKi, 1 mM, Millipore 688000), and 10 µg ml^-1^ human epidermal growth factor (Sigma Aldrich E9544). A series of filtration steps followed by passing the cell suspension through a 100, 70, and 40 µm cell strainer (Falcon 352360, 352350, 352340). An aliquot of filtered cell suspension was stained with Trypan blue, and the amount of live and dead cells was assessed. Cells were collected by centrifugation at 300xg for 7 minutes at 4°C and subjected to a dead cell removal assay (Dead cell removal Kit, Miltenyi Biotec 130-090-101) following the manufacturer protocol. In short, cells were suspended in 100 µl microbeads, MS columns (Miltenyi Biotec 130-042-201), 1.7 ml low protein binding tubes (Eppendorf, VWR 80077-230) and wide bore yellow pipette tips were used for the assay. The final cell pellet was kept at 0°C on ice and suspended in 50-100 µl A-Ri-Buffer (A-Ri-Buffer consisted of A-TE-2D supplemented with 20 µM ROCKi, and 100 ng ml^-1^ fibroblast growth factor 10 (FGF10, Preprotech 100-26) for a concentration of 1000-3000 cells per µl. A new cell aliquot was taken and assessed by the TC 20 automated cell counter (Bio-Rad 1450102) to obtain accurate cell number and viability of cells. Single preparations with a target cell recovery of 5000-6000 were loaded on a Chromium controller (10x Genomics, Single Cell 3’ v2 chemistry) to run single cell partitioning and barcoding utilizing the microfluidic system device. For next-generation sequencing of cDNA libraries the Illumina NextSeq500 System was applied.

Further, to improve the viability of primary human TE cells we developed a papain digestion assay and performed a different digestion protocol for experiments D36, D37, and D38. Cryopreserved distal fimbriae samples were thawed and washed as described above for the collagenase digestion protocol. Washed tissues were minced in 10 ml ice cold Wash Buffer and collected by centrifugation at 400xg for 6 minutes at 4°C. The tissue pellet was suspended in Papain Buffer (Papain Dissociation System, Worthington Biochemical LK003150; the Papain Buffer consisted of 1 vial Papain and 1 vial DNaseI dissolved in 5 ml Earle’s Balanced Salt Solution, the buffer was incubated for 20 minutes in a 37°C water-bath and adjusted to pH 7 by incubation at 5% CO_2_ in a 37°C incubator).

Tissue fragments in Papain Buffer were incubated for 2 hours in a 37°C water-bath. During incubation every 20 minutes the mixture was suspended by pipetting through a 5 ml pipette. The digestion was stopped by adding 15 ml Stop medium 1 (20% FBS in DMEM F12 50:50 (Corning 10-092-CV) and 0.1 mg ml^-1^ DNaseI (Millipore Sigma 10104159001) to the cell tissue mixture. Visible larger tissue pieces were allowed to settle for 5 minutes in the tube and the top cloudy cell suspension was added to a new tube. Cells were collected by centrifugation at 400xg for 7 minutes at 4°C, gentle suspended with 1.35 ml Albumin-ovomucoid inhibitor, prepared as to the kit’s protocol, and 3 ml Stop medium 1 was added. With the help of a pipette visible tissue pieces were removed from the cell solution. Cells were centrifuge as before, the cell pellet was gentle suspended by pipetting, 20 times, in 2 ml pre-warmed 37°C TryLE Express (Thermo Fisher 12604-013), 2 ml Stop medium 1 was added and the cell solution was filter through a 70 µm cell strainer followed by centrifugation as described above. Next, the pellet was resuspended with 3.5 ml Stop medium 1 and filtered through a 40 and 35 µm cell strainer followed by centrifugation at 400xg for 9 minutes at 4°C. The final cell pellet was dissolved using wide bore yellow pipette tips, in 50-100 µl A-Ri-Buffer, transferred to a 1.7 ml low protein binding tube and stored on ice. Cell counting, viability assessment and cDNA library preparations were done as for the D33, D34, and D35 experiments.

### Download and alignment of single-cell RNA sequencing data

For mouse samples, previously published sequencing data of mouse uterine tubes were downloaded from accession code GSE252786 in the Gene Expression Omnibus (GEO) database.

Human samples were sequenced and available as raw data files as mentioned previously. Additional raw sequencing files from human uterine tubes were acquired from Dinh et al (GSE151214) and Ulrich et al (GSE178101), which were both available in the GEO database ^14,15^. Samples acquired from Lengyel et al (EGAC00001003114) were made available through the European Genome-Phenome Archive (EGA) ^16^.

For sequence alignment, a custom reference for mice (mm39) and humans (GRCh38) was built using the cellranger (v.7.1.0, 10x Genomics) *mkref* function. The mm39.fa soft-masked assembly sequence and the mm39.ncbiRefSeq.gtf (release 109) genome annotation were used to form the custom reference for all of the mouse samples. The GRCh38.fa soft-masked assembly sequence and the GRCh38.108.gtf reference were the foundation for the custom human reference used to align all human samples. The raw sequencing reads were aligned to each species’ custom reference and quantified using the cellranger *count* function.

### Preprocessing and batch correction

All preprocessing was completed in R (v.4.2.0). For mouse samples, the datasets were preprocessed and corrected as previously described ^12^. The human samples were prepared identically beginning with ambient RNA removal. The output counts from cellranger alignment were modified using the *autoEstCont* and *adjustCounts* functions from SoupX (v.1.6.2) to generate a corrected counts matrix removing the ambient RNA signal (https://github.com/constantAmateur/SoupX). The corrected matrices were then processed individually following the standard pipeline from Seurat (v.4.3.0) using the *NormalizeData*, *FindVariableFeatures*, *ScaleData*, *RunPCA*, *FindNeighbors*, and *RunUMAP* functions (https://github.com/satijalab/seurat). The number of principal components used to construct a shared nearest-neighbor graph were chosen to account for 95% of the total variance for each individual dataset and subsequent combined datasets. To detect possible doublets, we used the package DoubletFinder (v.2.0.3) with inputs specific to each Seurat object. To establish a threshold for the proportion of artificial k nearest neighbors (pANN) values to distinguish between singlets and doublets, the estimated multiplet rates for each sample were calculated by interpolating between the target cell recovery values according to the 10x Chromium user manual. Homotypic doublets were identified using unannotated Seurat clusters in each dataset with the *modelHomotypic* function. After doublet detection calculations, the human and mouse datasets were merged separately. For mice, cells with greater than 30% mitochondrial genes, cells with fewer than 750 nCount RNA, and cells with fewer than 200 nFeature RNA were removed from the merged datasets. For humans, the mitochondrial gene threshold was set to 20% as more cells were available to account for a higher quality sample set. To correct for any batch defects between sample runs in each species, we used the harmony (v.0.1.0) integration method for each run rather than sample (https://github.com/immunogenomics/harmony).

### Clustering parameters and annotations

After merging the datasets and batch-correction, we employed Seurat’s pipeline to generate a combined dataset with dimensions reflecting 95% of the total variance were with a k.param of 70. Louvain clustering was then conducted using Seurat’s *FindClusters* with a resolution of 0.7 in both the human and mouse pooled datasets. The resulting 19 mouse clusters and 20 human clusters were annotated based on the expression of canonical genes and the results of differential gene expression (Wilcoxon Rank Sum test) analysis.

To better understand the epithelial populations, we subset epithelial cells from the distal region in mice and the fimbriae in humans. We then reclustered the 6 mouse epithelial populations and the 4 human epithelial populations. We reapplied harmony batch correction to each subset. For mice, the clustering parameters from *FindNeighbors* was a k.param of 50, and a resolution of 0.7 was used for *FindClusters*. The resulting 9 clusters within the mouse epithelial subset were further annotated using differential expression analysis and canonical markers. The Fibroblast-like population was suspected to be a doublet population as previously reported, and following analyses removed this population ^12^.

For the human fimbriae epithelial subset, we used a k.param of 70 and a resolution of 0.4 to generate cluster assignment for 9 epithelial clusters. However, three suspected doublet populations were removed that shared expression with immune, fibroblast, and endothelial populations respectively. A revised epithelial subset was reclustered with a k.param of 70 and a resolution of 0.7 to identify 9 clusters that all had unique expressions characterizing the fimbriae uterine tube epithelium, and these were used for later analyses.

### Pseudotime analysis

PHATE is dimensional reduction method to more accurately visualize continual progressions found in biological data ^27^. This was built on the phateR package (v.1.0.7) (https://github.com/KrishnaswamyLab/PHATE). Pseudotime values were calculated with Monocle3 (v.1.0.0) along the PHATE embeddings, which computes these trajectories with an origin set by the user (https://github.com/cole-trapnell-lab/monocle3) ^24–26^. The origin was set to be a progenitor cell state confirmed with lineage tracing experiments in mice. Human progenitors were identified based on shared expression of canonical genes in both mouse and human datasets. Pseudotime binning of each epithelial subset was completed by splitting the dataset into 40 equally sized bins stemming from the stem/progenitor state at pseudotime = 0.0. The bins were set to negative pseudotime scores for secretory lineages and positive scores for ciliated lineages. The average z-scored expression was calculated for each gene along the bins to identify genes that may reflect more subtle changes along trajectories. For gene signature scoring, we generated lists of genes from the Molecular Signatures Database (MSigDB) and used UCell (v.2.2.0) to score these gene signatures from lists across pseudotime bins or metadata features (https://github.com/carmonalab/UCell) ^40^.

### Mapping single-cell transcriptomes

All transcriptome mapping across species was completed using python (v.3.10.8). To explore conservation among the distal human and mouse datasets, we first loaded all human fimbriae and mouse distal filtered count matrices with scanpy (v.1.9.1). We then concatenated the sequencing files into a single dataset and subset them using annotations from the Seurat object for all the cells in the human and mouse datasets and the human and mouse epithelial subsets. We then converted these four datasets as self-assembling-manifold (SAM) objects using the sam-algorithm package (v.1.0.2) and followed the standard preprocessing steps (https://github.com/atarashansky/self-assembling-manifold) ^41^. Following SAM object preparation, we then applied the SAMap (v.1.0.12) pipeline to SAM objects of all cells and epithelial subsets to map both mouse and human datasets together (https://github.com/atarashansky/SAMap) ^21^.

### Conserved and analogous feature selection

To calculate the top correlated features within the combined epithelial SAMap object, we first calculated the gene pairs and aligned through a correlation of 0.2 using the *GenePairFinder* and *find_all* functions. The identified markers were used to find orthologous genes for conserved lineage characterization.

To identify orthologous genes that were highly correlated but also species specific, we began with exporting Wilcoxon Rank Sum test results from Seurat’s *FindAllMarkers* function on epithelial subsets for mice and humans. We then converted the top distinguishing genes for each epithelial cell state in both species to their respective ortholog. These gene lists were paired and read through SAMap’s *query_gene_pair* function with gene-gene correlation as the output. The results were reorganized to display the top conserved gene pairs as marked by the highest correlation and species-specific gene pairs as marked by the lowest correlation scores.

To identify analogous genes, the top correlated gene pairs found using the *GenePairFinder* and *find_all* functions were annotated specifically to highlight surface markers for the human genes. The gene pairs with the human gene annotated as a surface marker were retained, while all other gene pairs were filtered out. Since SAMap relies on sequence similarity from BLAST score results, analogous terms were defined by their inverse BLAST score. The terms with the highest correlation score (>0.4) and inverse BLAST scores (>0.010) were subjected to additional filtering. The next filtering measure was based on species specificity. The fold change of average expression from progenitors compared to other cell states were calculated from pseudotime lineage tracing of human epithelial cells. The top candidates were ranked and selection from the top 10 terms was based on biological interest for further validation studies.

### Adherent primary human TE culture

Cryopreserved fimbriae samples were thawed and digested with collagenase type I as described above for D33, D34, and D35 for 45 minutes at 37°C, cells were collected by centrifugation. The cell/tissue pellet was suspended with 9 volumes Stop medium 2 (5% FBS in Basic Medium which consisted of DMEM F12 50:50, 1% GlutaMax-I, PS, 1 mM sodium pyruvate (100 mM, Thermo Fisher 11360-070), 1% MEM Non-Essential Amino Acids (100x, MEM NEAA, Thermo Fisher 11140-050) and centrifuged at 400xg for 6 minutes at 4°C. Unless otherwise noted all following centrifugations were carried out at this setting. Harvested cells were suspended in 2 ml TrypLE Express, incubated for 4 minutes in a 37°C water bath, 18 ml Stop medium 2 was added and the cells were collected by centrifugation. Cell pellets were suspended in A-TE-2D modified medium (A-TE-2D modified was made up of 2.5% FBS, Basic medium, 25% Wnt-3A/R-Spondin1/Noggin conditioned medium (L-WRN cells, ATCC CRL-3276), 10 µM ROCKi, 10 µg ml^-1^ EGF, 1 mM nicotinamide (Sigma Aldrich A9165), 1 mM N-acetyl-L-cysteine (Sigma Aldrich A9165), 10 µM p38 inhibitor (p38i, SB202190, Sigma Aldrich S7067), 2.5 µM TGF-ß RI Kinase Inhibitor VI (Ti, SB431542, ALK4/5, Millipore 616464)), and live and dead cells were counted after Tryphan blue staining. Adherent cultures of the isolated cells 0.5 x 10^6^ cells in 10 ml per gelatinized 10 cm cell culture dish were started. Small 1 mm epithelial tissue clusters were added and cultivated in the same dishes. Larger tissue pieces were discarded. Primary TE cells grew 7 to 11 days with medium changes every 3 to 4 days.

### Cell population separation with magnetic beads (MACS)

Adherent cultivated human primary TE cells were harvested after treatment with 1.5 ml TrypLE Express per 10 cm dish, digestion was ended by adding Stop medium 2, and cells were filtered through a 40 µm mesh. Total cells were resuspended in 3 ml Stop medium 2, passed through a 35 µm cell strainer cap into a 5 ml polystyrene tube (Falcon REF352235), an aliquot was stained with Tryphan blue, cells were counted and centrifuged at 400xg for 7 minutes at 4°C. All further centrifugations were done with this setting. If not mentioned otherwise for intermediate steps cells and buffers were kept at 0°C on ice. Total cells at a concentration of 5x 10^6^ cells ml^-1^ were suspended in A-MACS Medium (Basic Medium with the addition of 2% B27 (Thermo Fisher 175-04-044), 1% N2 (Thermo Fisher 17502-048), 1 mM nicotinamide, 1 mM n-acetyl-L-cysteine, 10 µM ROCKi, 10 µM p38i). We targeted four lineage negative cell types (Lin^-^) with specific antibodies. Endothelial cells (CD31, eBioscience 13-0319-82, biotin), T cells (CD45R, eBioscience 13-0452-81, biotin), Macrophages (CD11b, eBioscience 13-0118-82, biotin), and Erythrocytes (CD235a, eBioscience 13-9987-82, biotin). All four Lin^-^ antibodies were added at a concentration of 3 µg each for 10^6^ suspended target cells and incubated for 25 minutes at 0°C, with every 5 minutes gentle shaking of the solution. After the incubation 1.5 ml of MACS Buffer 2 (0.1% bovine serum albumin (BSA, Sigma Aldrich A9647) in PBS, and 2 mM ethylenediaminetetraacetic acid (EDTA, pH 7)) was added and the suspension centrifuged. Pelleted cells were suspended in 250 µl MACS Buffer 2 , transferred to a 1.7 ml low protein binding tube, 150 µl of washed magnetic beads (Dynabeads Biotin Binder, Thermo Fisher 11047) in MACS Buffer 1 (0.1% BSA in PBS) were added and the mixture was incubated for 20 minutes at 4°C with constant rotation to facilitate binding of Lin^-^ cells to the magnetic beads. Washing of beads was accomplished in advance by suspending the beads in the original company vial, transferring 150 µl of beads to a 1.7 ml low attachment tube, adding and suspending them in 1 ml MACS Buffer 1, placing the tube in a magnet rack for 2 minutes at 25°C, with help of a yellow pipette tip taking out the supernatant, removing the tube from the magnet rack, and suspending the washed beads with the same volume of MACS buffer 1 as the initial volume taken from the beads vial, 150 µl. Following the binding of Lin^-^ cells to the beads, the tube was placed in a magnet rack for 2 minutes at 25°C, the supernatant containing cells cleaned from the Lin^-^ population, was carefully transferred to a new 1.7 ml tube. Beads were discarded. The cell supernatant was centrifuged. Pelleted cells were resuspended with 300 µl A-MACS Medium, the cell concentration was assessed, the volume was divided into two tubes. To 150 µl PLA2R1 antibody (Novus Biotechne, NBP2-50248B, biotin) was added at a concentration of 6 µg per 10^6^ cells, and to the second tube, 150 µl, ROBO1 antibody (Novus Biotechne, FAB71181B, biotin) was added at an equal concentration. Both tubes were placed on ice for 25 minutes to incubate, then 1 ml MACS Buffer 2 was added as a wash step. Cells were collected by centrifugation. Each pellet was resuspended in 150 µl MACS buffer 2, 100 µl washed beads were added, the mixture was placed on a rotator and incubated for 20 minutes at 4°C. Beads and unbound cells were separated by placing the tubes in a magnet rack for 2 minutes. The supernatant from each marker tube e.g. the PLA2R1 negative unbound cell population and ROBO1 negative unbound cells were transferred into a new 1.7 ml tube and kept on ice. Tubes containing the beads with attached PLA2R1 positive cells and ROBO1 positive cells were taken out from the magnetic rack. Each bead pellet was suspended with at 37°C pre-warmed 200 µl Release buffer containing 1 Unit DNase per µl (Release Buffer, 1% FBS in Basic Medium, with 1 mM CaCl_2_, 5 mM MgCl_2_, and 1U/µl DNase I) (DNase I, Worthington LS002006), then incubated for 10 minutes in a 37°C water-bath, beads were once mixed by pipetting then collected with a magnet. The supernatant, released marker labeled cells, were collected and stored on ice. A second releasing procedure followed. Released cell solutions were pooled 900 µl A-MACS Medium was added and the cells collected by centrifugation. Cell pellets were suspended in 100-150 µl GyneCult medium (GC, StemCell Technologies 100-1247).

### Human TE organoid cultivation

MACS isolated cell subpopulations, negative and positive marker fractions (1-5x 10^4^ cells per rim assay), three assays per population were prepared as previously described ^38^. In short, cells suspended in GC Medium were mixed with Matrigel (Corning 356231) in the ratio of 30:70. The mixture was spread around the rim of an uncoated 24 well plate. After incubation at 20 minutes at 37°C in a 5% CO_2_ incubator 500 µl GC medium was added to the middle of the well. Every 3 to 4 days the media was changed. Organoid rim assays were photographed after 12 days. Afterwards, fixed for histological preparation.

### Organoid histological preparation and analysis

At 25°C the GC medium was removed from each rim assay without disturbing the Matrigel organoid mixture, the plate was placed at 0°C on ice, and 1 ml Fixation Buffer (4% Paraformaldehyde in 500 mM PIPES (Bioworld 41620140-2), 25 mM MgCl_2_, 50 mM EDTA) was added to the middle of a 24 well. Organoids were fixed for 3.5 hours on ice. Then, the Fixation Buffer was taken out from each well, the plate moved to 25°C, 0.5 ml IF Buffer (0.2% Triton, 0.05% Tween in PBS) was added, with help of a wide bore yellow tip organoids were suspended in IF Buffer and transferred to a 1.7 ml tube. An additional 0.5 ml IF Buffer was used to collect all remaining organoids from the well and combined with the first IF Buffer solution. Organoids were collected by centrifugation at 300xg for 5 minutes at 25°C. The organoid pellet was washed 3 times with 600 µl PBS. The final organoid pellet was suspended in 600 µl 70% ethanol and stored at 4°C overnight. The process was continued by pelleting the organoids and suspending them in 100 µl of melted 60°C Histogel (Thermo Fisher REF HG-4000-012). The mixture was placed on a Parafilm lined petri dish, the drop was solidified for 10 minutes at 4°C, transferred to a histological plastic cassette lined with foam pads, numbered and stored in 70% ethanol until paraffin embedding.

### Organoid analysis

Images of each well (technical replicate) from three separate experiments (biological replicates) were quantified for the organoid diameters of PLA2R1-bound, PLA2R1-unbound, ROBO1-bound, and ROBO1-unbound groups. The diameters were measured using Fiji (National Institutes of Health, Bethesda, MD, USA).

For differentiation marker investigation, 4 μm-thick tissue sections were collected from prepared paraffin-embedded blocks of organoids. The organoids were stained for secretory marker (KRT7) and ciliated marker (FOXJ1). Images of organoids were collected on a 40x objective with a Zeiss LSM 710 Confocal Microscope using the tile scan setting with a pixel dwell of 1.58 μsec and an averaging of 2 using the ZEN (blue edition, Zeiss) software. Organoids were quantified for their abundance of nuclei, KRT7+ cells, and FOXJ1+ cells. The percent was calculated by the abundance of the respective differentiation markers divided by the total nuclei in each organoid.

### Experimental animals

The Tg(KRT5-cre/ERT2)2Ipc/Jeldj (K5-Cre-ERT2, Stock number 018394), and *Gt(ROSA)26Sor^tm9(CAG-tdTomato)Hze^* (*Rosa-loxp-stop-loxp-*tdTomato/Ai9, Stock number 007909), mice were obtained from The Jackson Laboratory (Bar Harbor, ME, USA). The *Trp53^loxP/loxP^* and *Rb1^loxP/loxP^* mice, which have *Trp53* and *Rb1* genes flanked by *loxP* alleles, respectively, were a gift from Dr. Anton Berns. Mice were bred on a mixed FVB/N and BL6/J background. Their genotype was confirmed by PCR amplification (Supplementary Table 2). Only females were used for our studies of the female reproductive tract. The number of animals used in every experiment is indicated as biological replicates in figure legends and supplementary tables. All the experiments and maintenance of the mice followed ethical regulations for animal testing and research. They were approved by the Cornell University Institutional Laboratory Animal Use and Care Committee (protocols numbers 2000-0116 and 2001-0072). Mice were housed within a 10/14 light cycle. The lights came on at 5 a.m. and went off at 7 p.m., the humidity ranged from 30-70 %, and the ambient temperature was kept at 72°F +/- 2°F.

### Tamoxifen induction

For trauma induction assays, 6-10-week-old K5-Cre-ERT2 Ai9 and K5-Cre-ERT2 *Trp53^loxP/loxP^ Rb1^loxP/loxP^* Ai9 mice of the same age received a single dose of tamoxifen (100 µg g^-1^ body weight, 8 µl g^-1^ body weight, 12.5 mg ml^-1^ in corn oil; Sigma-Aldrich, T5648) by intraperitoneal (i.p.) injection. Tamoxifen was injected immediately after the trauma surgery.

### Microsurgical uterine tube injury

Mice at diestrus stage underwent surgery. Surgeries on experimental animals were performed as previously described with specific modifications ^42^. After the reproductive tract was exposed, an incision in the bursa above the infundibulum was made, using blunt fine forceps the infundibulum was grasped and squeezed for 30 seconds. Next the reproductive tract was placed back into the peritoneum following the previously described surgical procedure ^42^. The undamaged contra-lateral oviduct served as control tissue. After the surgery, mice were immediately injected with tamoxifen. K5-Cre-ERT2 Ai9 mice were collected at chase point 1, 4, 10, and 30 days. K5-Cre-ERT2 *Trp53^loxP/loxP^ Rb1^loxP/loxP^* Ai9 mice were euthanized at the same chase points and upon sickness between 171 – 208 days of age.

### Immunohistochemistry and image analysis

For all mouse experiments, collected tissues were fixed with 4% paraformaldehyde overnight at 4°C followed by standard tissue processing, paraffin embedding, and preparation of 4 μm-thick tissue sections. For immunohistochemistry, antigen retrieval was performed by incubation of deparaffinized and rehydrated tissue sections in boiling 10 mM sodium citrate buffer (pH 6.0) for 15 min. The primary antibodies for fluorescently labeled immunohistochemistry were incubated at room temperature for 1 hour prior to incubation with secondary fluorescent antibodies (dilution 1:200, 1 hour, at room temperature, RT). Vectashield mounting medium with DAPI (Vector Laboratories, Burlingame, CA, USA; H-1800) was used as the counterstain for nuclei detection. Otherwise, primary antibodies were incubated at 4 °C overnight, followed by incubation with secondary biotinylated antibodies (dilution 1:200, 1 hour, at room temperature, RT). Modified Elite avidin-biotin peroxidase (ABC) technique (Vector Laboratories, Burlingame, CA, USA; pk-6100) was performed at room temperature for 30 min. Hematoxylin was used as the counterstain. All primary and secondary antibodies used for immunostaining are listed in Supplementary Table 3.

For quantitative studies, sections were scanned with a ScanScope CS2 (Leica Biosystems, Vista, CA), 40x objective, a Zeiss LSM 710 Confocal Microscope for tile scans with a pixel dwell of 1.58 μsec and an averaging of 2 using the ZEN (blue edition, Zeiss) software, or a Leica TCS SP5 Confocal Microscope, 40x objective, followed by the analysis with Fiji software (National Institutes of Health, Bethesda, MD, USA).

### Pathological evaluation

All mice in carcinogenesis experiments were subjected to gross pathology evaluation during necropsy. Potential sites of ovarian carcinoma spreading were prioritized, such as the omentum, regional lymph nodes, uterus, liver, lung, and mesentery. In addition to the uterine tube and ovary, pathologically altered organs, as well as representative specimens of the brain, lung, liver, kidney, spleen, pancreas and intestine, intra-abdominal lymph nodes, omentum and uterus were fixed in 4% PBS buffered paraformaldehyde. They were then evaluated by microscopic analysis of serial paraffin sections stained with hematoxylin and eosin and subjected to necessary immunostainings. All early atypical lesions were diagnosed based on their morphology, staining for tdTomato, extent of proliferation, and detection of deleted (floxed-out) *Trp53* and *Rb1* as described earlier ^43^. STICs were identified based on pronounced cellular atypia, the loss of cell polarity, high proliferative index, and *Trp53* deletion. The locations of all lesions were determined by three-dimensional reconstruction of 4-mm-thick serial sections as described previously ^44^.

### Statistical analyses

Statistical comparisons were performed using a two-tailed unpaired *t* test and Analysis of Variance (ANOVA) with Tukey-Kramer Multiple Comparisons Test with InStat 3 and Prism 10.6.1 software (GraphPad Software Inc., La Jolla, CA, USA).

## Data Availability

The raw sequencing data generated in this study have been deposited in the Gene Expression Omnibus (GEO) database under accession code GSE324855. Additional human datasets were acquitted from Dinh et al (GSE151214), Ulrich et al (GSE178101), and Lengyel et al (EGAC00001003114). All processed Seurat objects from mouse scRNA-seq experiments are available in the Dryad repository at https://doi.org/10.5061/dryad.4mw6m90hm. All processed Seurat objects from the human dataset and all SAMap objects are available in the Dryad repository after manuscript acceptance. Supplementary marker identification and correlation analysis for mouse, human, and combined datasets are available on GitHub (https://github.com/coulterr24/Uterine_Tube_Comparitive_Atlas). All data generated in this study are provided in the Source Data file. The remaining data are available within the Article, Supplementary or Source Data file.

## Code Availability

All code for data preprocessing and figure generation of scRNA-seq data are available through GitHub (https://github.com/coulterr24/Uterine_Tube_Comparitive_Atlas).

## Acknowledgements

We thank Peter A. Schweitzer, Director of the Cornell Genomics Facility for his invaluable assistance with single-cell RNA sequencing. This work has been supported by NIH grants (CA182413, CA260115 and CA248524 to AYN, and U54AG079779 to BDC), Ovarian Cancer Research Fund grant (327516) to AYN, the Chan Zuckerberg Initiative Single-Cell Biology Data Insights grant (DAF2023-323354) to BDC, the Kansas Institute for Precision Medicine via the NIGMS (P20 GM130423), the KU Cancer Center’s Cancer Center Support Grant (P30 CA168524), and the Honorable Tina Brozman Foundation, Inc. to AKG, the NSF Graduate Research Fellowship Program (GRFP) awarded to CQR (DGE-2139899) and NIH 1S10RR025502 grant to the Cornell Institute of Biotechnology Imaging Facility for the Zeiss LSM 710 Confocal Microscope.

## Author contributions

CQR, AFN, and AYN designed experiments; CQR, DJF, CSA, BAH, MMH, DKW, and AFN performed experiments; CQR, DKW, and BDC carried out bioinformatics analyses; DJF, AY, ES, DM and AYN performed pathological evaluation; ES, AY, AKG, and DM provided resources; CQR, AFN and AYN wrote the paper.

