## Supplementary Information for "Comparative single-cell atlases reveal injury-driven tubal epithelial regeneration as a window for ovarian carcinoma initiation"

Ralston et al.

### Supplementary Figures

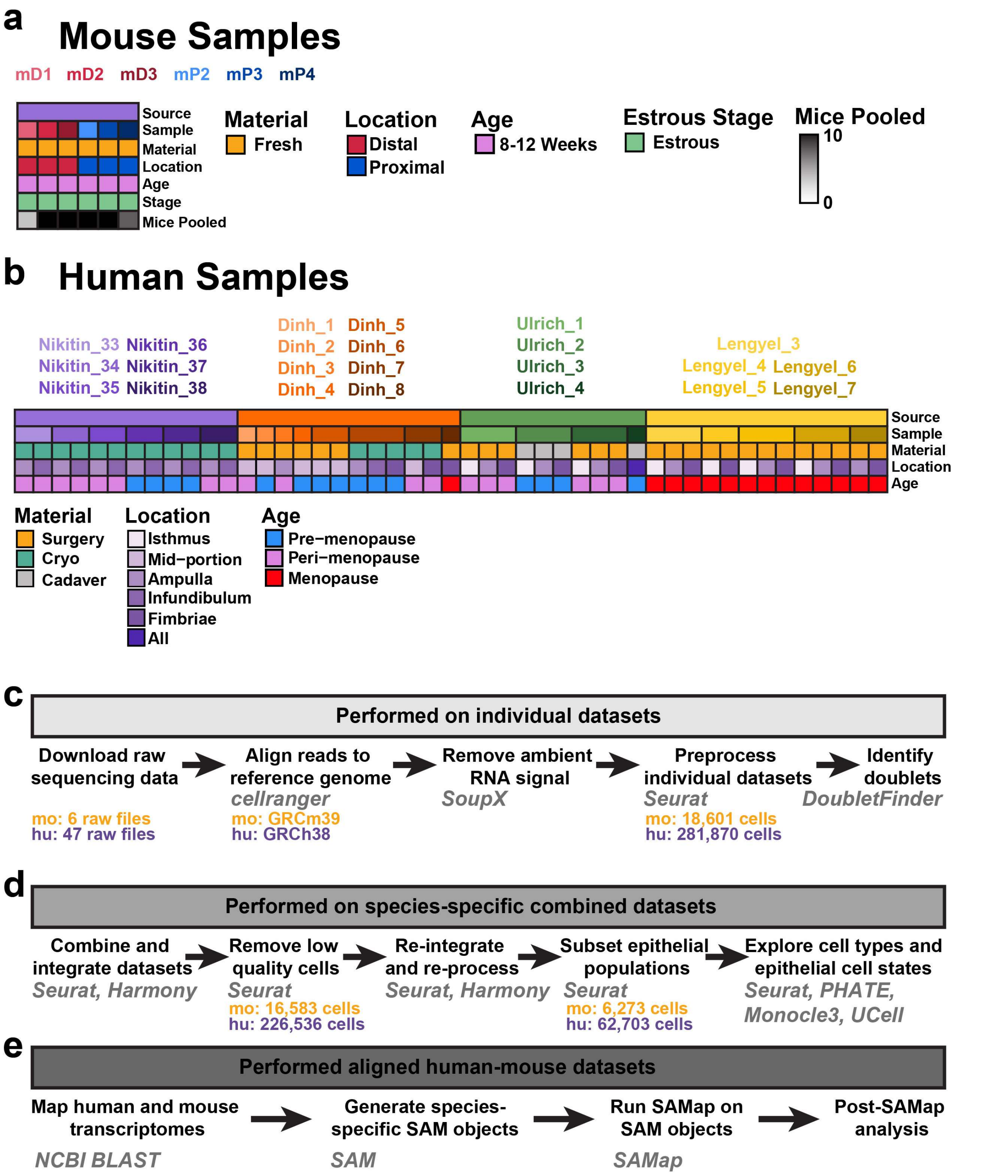


**Supplementary Figure 1. Sample preparation and preprocessing.** (**a**) Metadata visualization for the mouse scRNA-seq dataset. (**b**) Metadata visualization for the human scRNA-seq dataset. (**c-e**) Dataset preparation workflow beginning with individual dataset preparation (c), combined dataset batch correction and exploration (d), and multiple species alignment using SAMap.

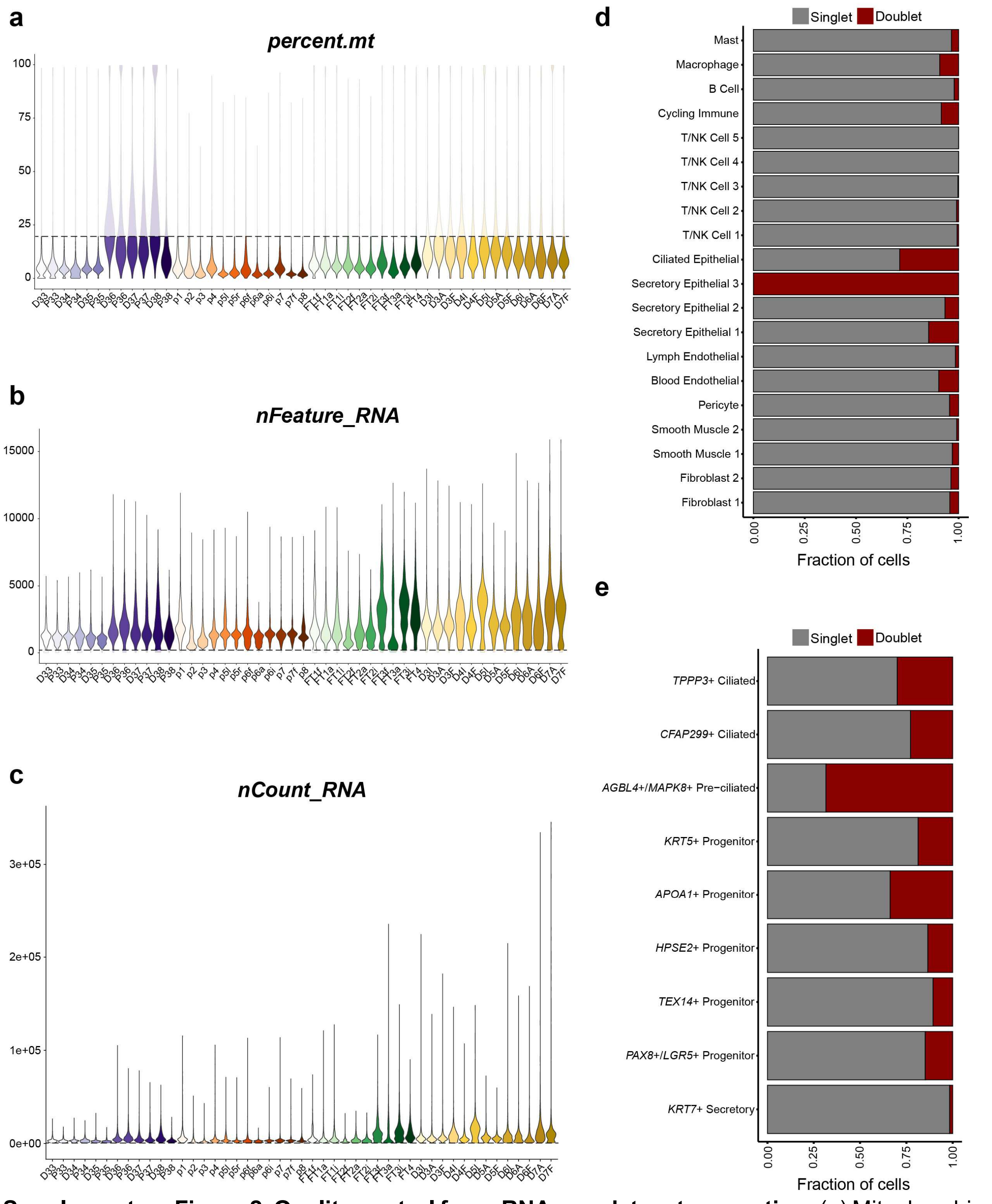


**Supplementary Figure 2. Quality control for scRNA-seq dataset generation.** (**a**) Mitochondria percentage was used to filtered cells with values greater than 20%. (**b**) RNA feature counts filtered out cells with values fewer than 200. (**c**) RNA counts filtered out cells with values fewer than 750. (**d-e**) Doublet detection using DoubletFinder for visualization of possible doublet populations in the whole human (d) and distal epithelial human (e) datasets.

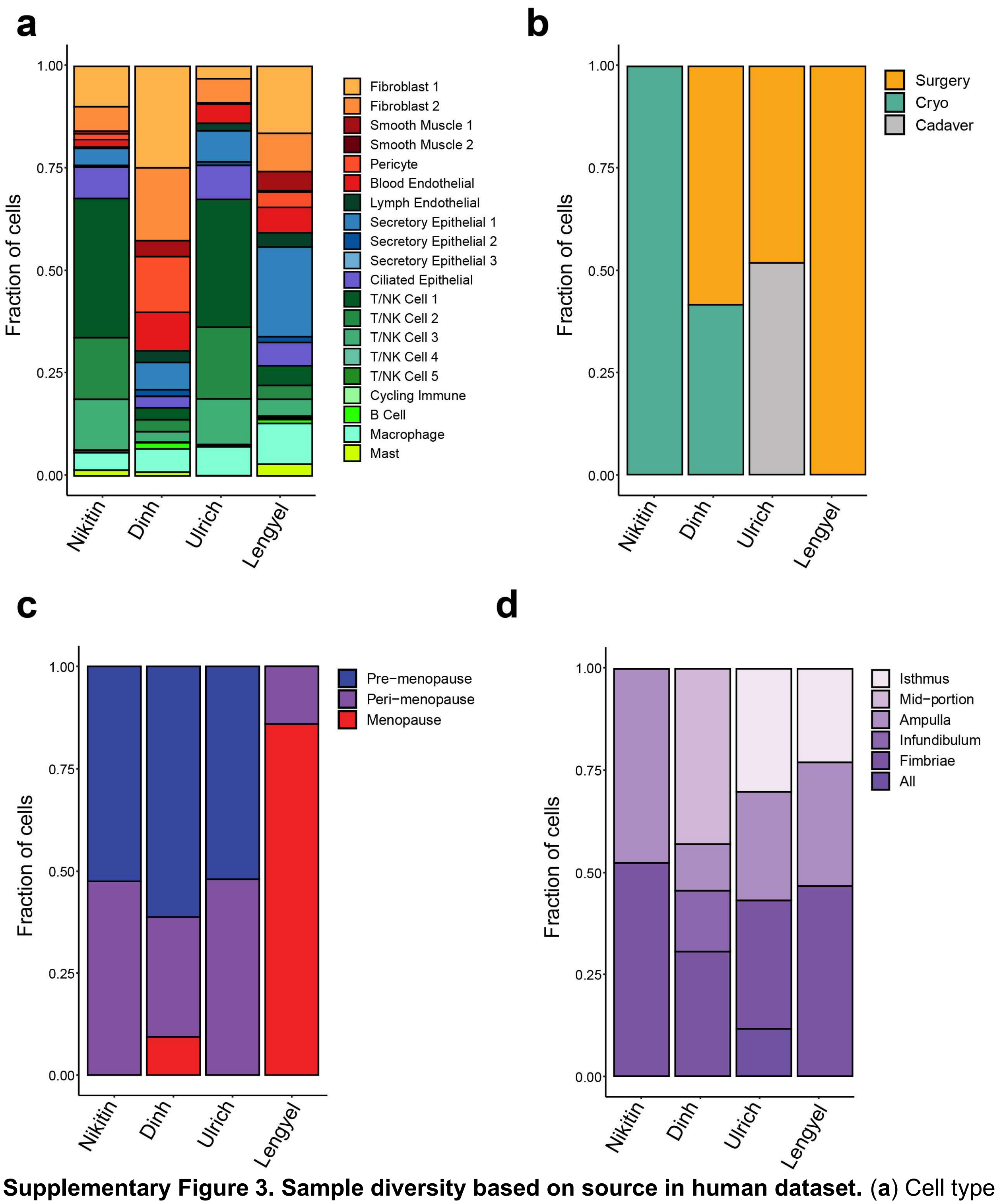


**Supplementary Figure 3. Sample diversity based on source in human dataset.** (**a**) Cell type distribution by source in which the sequencing files were obtained (**b**) Source differences in collected materials within the dataset. (**c**) Age group differences based on the source from which samples were derived. (**d**) Source-based distribution of regions collected for the human dataset.

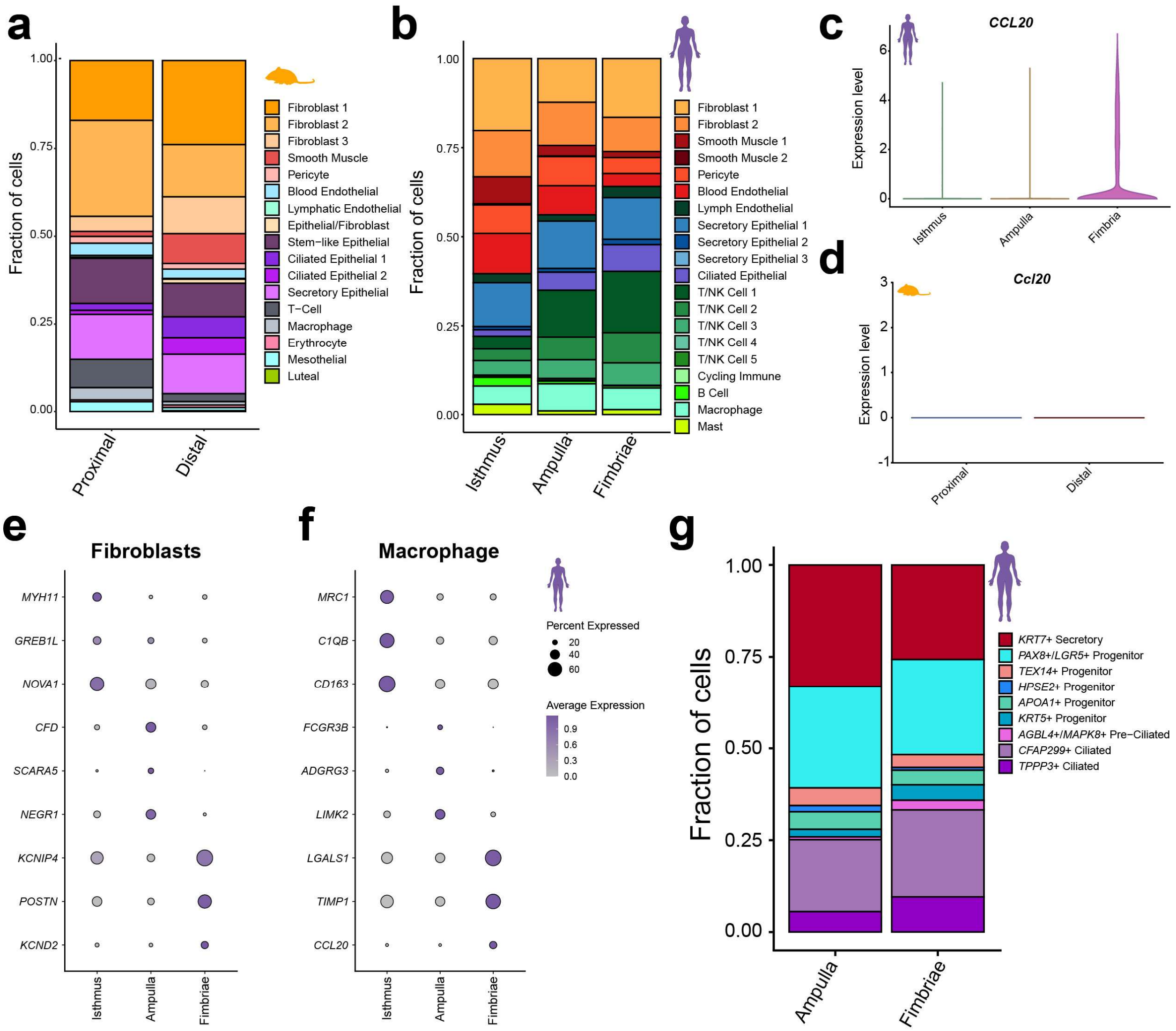


**Supplementary Figure 4. Regionally-associated differences in human and mouse datasets.** (**a**) Stacked bar chart showing cellular distribution for distal and proximal regions from the mouse uterine tube. (**b**) Stacked bar chart showing cellular distribution for isthmus, ampulla, and fimbria regions from the human uterine tube. (**c-d**) Violin plot expression of *CCL20* split across regions for macrophages from human (c) and mouse (d) datasets. (**e-f**) Top distinguishing markers exclusive to regions for human fibroblasts (e) and macrophages (f). (**g**) Stacked bar plot for cellular distribution among the ampulla and fimbria regions in human distal epithelial subsets.

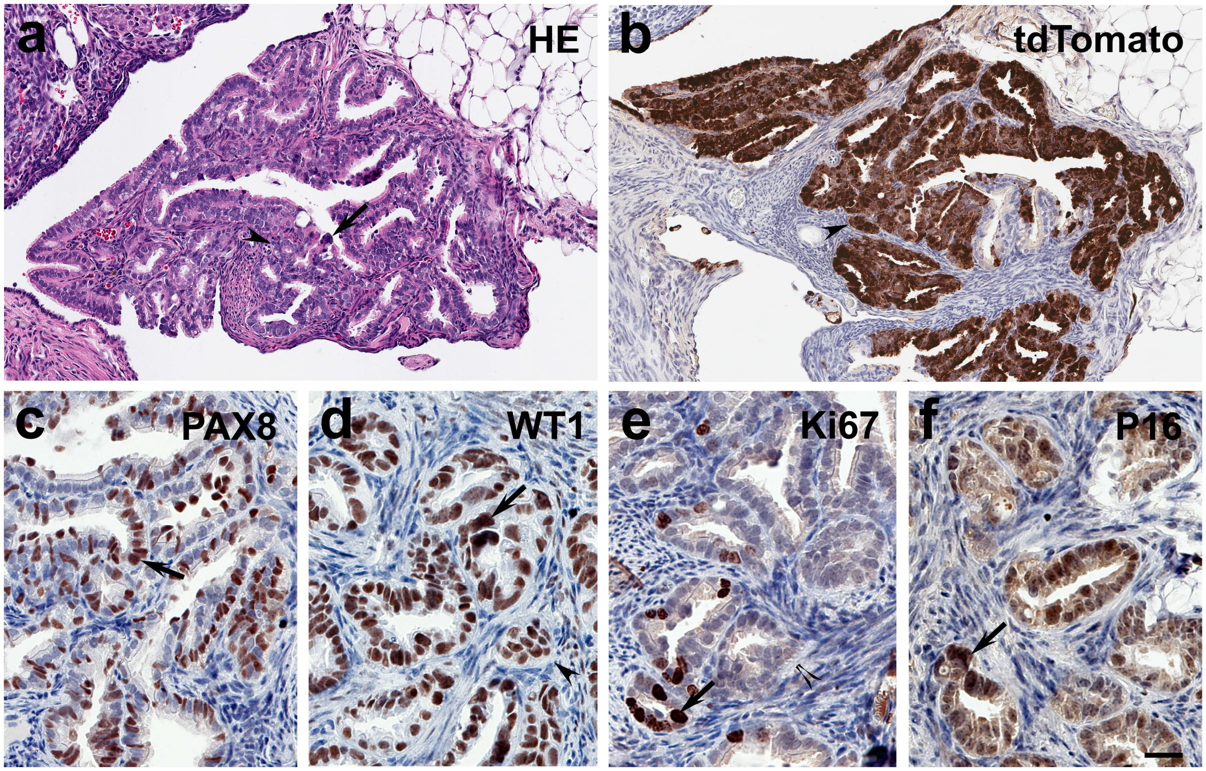


**Supplementary Figure 5. Advanced neoplastic lesions in injured uterine tube of Krt5-CreERT *Trp53*$^{loxP/loxP}$ *Rb1*$^{loxP/loxP}$ Ai9 mice.** (**a-f**) Lesions are marked by pronounced nuclear atypia (arrows) and stromal invasion (arrowheads), and expression of PAX8 (c), Wilms tumor 1 (WT1, d), Ki67 (e) and P16 (f). Hematoxylin and eosin (HE, a) staining, and immunostainings (brown color) for tdTomato, PAX8, WT1, Ki67 and P16 by Elite ABC method, hematoxylin counterstaining. Scale bar, 50 µm (a, b) and 25 µm (c -f).

### Supplementary Tables

**Supplementary Table 1.** Neoplastic TE lesions in injured and non-injured contralateral uterine tubes after Cre-*LoxP* mediated inactivation of *Trp53* and *Rb1* in *Krt5*+ pre-ciliated cells

| Injury | Yes | No |
|---|---|---|
| Cases, N | 18 | 18 |
| First detection (DPI) | 26 | 120 |
| End point (DPI) | 208 | 208 |
| Targeted cells (%)[#] | 0.83±0.49 | 0.83±0.49 |
| TE lesions (%)* | 100 | 44 |
| STIC | 8 | 6 |
| HGSC[&] | 10 | 2 |

[#] According to tdTomato detection in Ai9 mice crossed to "Cre" mice

Mean ± SD

*TE lesions: injured vs non injured Fisher's exact P = 0.0003

[&]HGSC: Fisher's exact P = 0. 0116

**Supplementary Table 2.** List of genotyping primers

| **Mouse strain*** | **Gene detected** | **Primer name** | **Sequence (5' → 3')** | **PCR products** |
|---|---|---|---|---|
| Rosa-loxP-stop-loxP-tdTomato/Ai9 | Ai9 | Ai9-1 | AAGGGAGCTGCAGTGGAGTA | 297bp-Wild type |
| | | Ai9-2 | CCGAAAATCTGTGGGAAGTC | 196bp-Mutant |
| | | Ai9-3 | GGCATTAAAGCAGCGTATCC | |
| | | Ai9-4 | CTGTTCCTGTACGGCATGG | |
| K5-Cre-ERT2 | *Cre* | Cre5' | GGACATGTTCAGGGATCGCCAGGC | 269bp-Mutant |
| | | Cre3' | GCATAACCAGTGAAACAGCATTGCT | |
| *Rb1*$^{loxP/loxP}$ | *Rb1* | Rb212 | CGAAAGGAAAGTCAGGGACATTGGG | 295bp-Floxed |
| | | Rb183' | GGAATTCCGGCGTGTGCCATCAATG | 247bp-Wild type |
| | | Rb19E | AGCTCTCAAGAGCTCAGACTCATGG | 269bp-Knockout |
| *Trp53*$^{loxP/loxP}$ | *Trp53* | P531F5' | GTGCCCTCCGTCCTTTTTCGCAATC | 316bp-Floxed |
| | | P5310F5' | GTTAAGGGGTATGAGGGACAAGGTA | 163bp-Wild type |
| | | P53102.3' | CCATGAGACAGGGTCTTGCTATTGT | 198bp-Knockout |

*See Methods for strain descriptions.

**Supplementary Table 3.** List of antibodies used for immunostaining

| Antigen, conjugation | Antibody source, catalogue number | Clone | Host | Retrieval | Dilution |
|---|---|---|---|---|---|
| FoxJ1 | Novus Biologicals, AF3619-SP | PC[#] | Goat | Citrate | 1:800 (IF*) |
| Ki67 | Thermo Fisher, 14-5698-82 | SolA-15 | Rat | Citrate | 1:4000 (IHC[&]) |
| KRT7 | Novus Biologicals, NBP2-44814 | OV-TL12/30 | Mouse | Citrate | 1:200 (IF) |
| P16 | Abcam, Ab241543 | PABLO-33B | Rat | Citrate | 1:500 (IHC) |
| p73 | Abcam, Ab40658 | EP436Y | Rabbit | Citrate | 1:200 (IF) |
| PAX8 | Proteintech, 10336-1-AP | PC | Rabbit | Citrate | 1:4000 (IHC), 1:400 (IF) |
| RFP/tdTomato | Rockland Immunochemical, 600-401-379S | PC | Rabbit | | 1:4000 (IHC) |
| Periostin | Abcam, Ab14041 | PC | Rabbit | | 1:3000 (IHC) |
| PLA2R1 | Novus Biologicals, NBP1-84449 | PC | Rabbit | Citrate | 1:200 (IF) |
| ROBO1 | Novus Biologicals, NB600-1254 | PC | Rabbit | Citrate | 1:200 (IF) |
| Wilm's Tumor Protein 1 | Abcam, Ab267377 | EPR-23963 | Rabbit | Citrate | 1:500 (IHC) |
| Anti-goat IgG, Alexa Fluor 488 | Invitrogen, A-11055 | PC | Donkey | | 1:200 |
| Anti-mouse IgG, Alexa Fluor 488 | Thermo Fisher, A-11001 | PC | Goat | | 1:200 |
| Anti-rabbit IgG, biotinylated | Vector Labs, BA-1000-1.5 | PC | Goat | | 1:200 |
| Anti-rabbit IgG, Alexa Fluor 594 | Thermo Fisher, A-21207 | PC | Donkey | | 1:200 |
| Anti-rat IgG, biotinylated | Vector Labs, BA-4000-1.5 | PC | Rabbit | | 1:200 |

*IF, Immunofluorescence; [#]PC: Polyclonal; [&]IHC, Immunohistochemistry (ABC eliite method)
